# MUC17 is an essential small intestinal glycocalyx component that is disrupted in Crohn’s disease

**DOI:** 10.1101/2024.02.08.578867

**Authors:** Elena Layunta, Sofia Jäverfelt, Fleur C. van de Koolwijk, Molly Sivertsson, Brendan Dolan, Liisa Arike, Sara Thulin, Bruce A. Vallance, Thaher Pelaseyed

## Abstract

Crohn’s disease (CD) is the chronic inflammation of the ileum and colon triggered by bacteria, but insights into molecular perturbations at the bacteria-epithelium interface are limited. We report that membrane mucin MUC17 protects small intestinal enterocytes against commensal and pathogenic bacteria. In non-inflamed CD ileum, reduced MUC17 levels correlated with a compromised glycocalyx, allowing bacterial contact with enterocytes. *Muc17* deletion in mice rendered the small intestine prone to atypical infection while maintaining resistance to colitis. The loss of Muc17 resulted in spontaneous deterioration of epithelial homeostasis and extra-intestinal translocation of bacteria. Finally, Muc17-deficient mice harbored specific small intestinal bacterial taxa observed in CD. Our findings highlight MUC17 as an essential line of defense in the small intestine with relevance for early epithelial defects in CD.

**One-sentence summary:** Membrane mucin MUC17 protects enterocytes against bacterial attachment and constitutes an early defect in Crohn’s disease.

## Introduction

Mucins are critical for the gastrointestinal defense system (*1, 2*). In the mouse distal colon, the gel-forming Muc2 forms an attached impenetrable mucus layer that separates bacteria from the epithelium (*3*). Defects in core mucus components constitute early steps in the pathogenesis of ulcerative colitis, a form of inflammatory bowel disease (IBD) in humans (*4*). In contrast, the mucus of the small intestine is non-attached and permeable to luminal material including bacteria (*5*). This regiospecific difference demands a distinct defense mechanism to manage host-microbiota interactions. In this context, the role of membrane mucins in small intestinal enterocytes is unexplored, due to limited insight into their gene sequences (*6*) and difficulties preserving them in fixed tissues. MUC17, alongside MUC13, is a conserved heavily *O*-glycosylated membrane mucin positioned at the apical brush border of differentiated enterocytes (*7, 8*) (Fig. 1A). In the mouse small intestine, the homeostatic cytokine IL-22 induces Muc17 expression and the establishment of a surface glycocalyx that in *ex vivo* experiments prevents direct attachment of bacteria to the brush border (*9*).

**Fig. 1.**
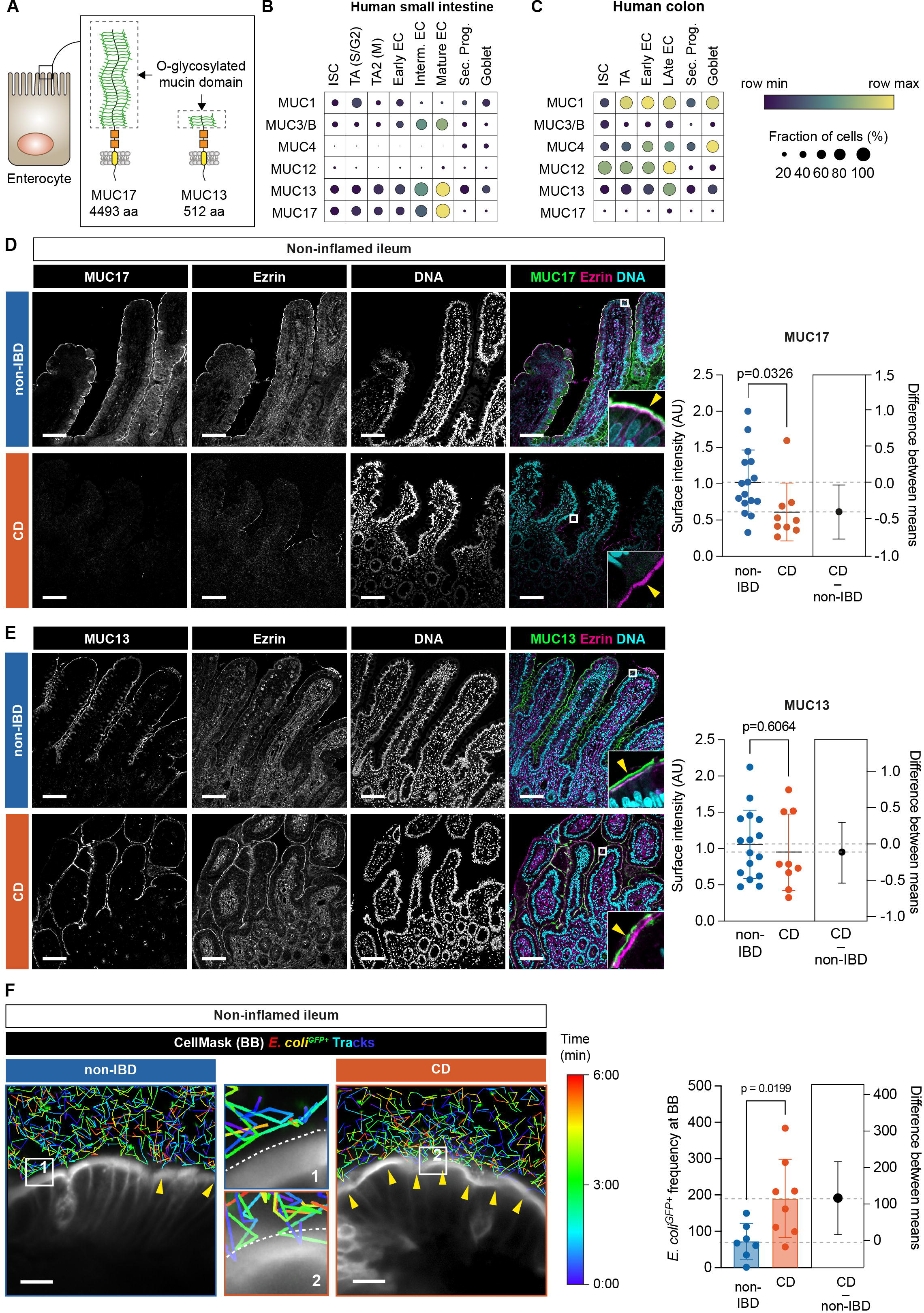
Non-inflamed ileum of CD patients expresses reduced levels of MUC17 and carries a glycocalyx permeable to luminal bacteria. (**A**) Schematic illustration of human MUC17 and MUC13 position at the enterocyte brush border. Dashed boxes show the *O*-glycosylated mucin domain in each mucin. aa, amino acids. (**B**) Dot plot showing mRNA expression of membrane mucin genes at single-cell resolution in epithelial cell types of the human small intestine. EC, enterocyte; ISC, intestinal stem cell; TA, transit amplifying; Sec. Prog., secretory progenitor; Goblet, goblet cell. S/G2, S- and G2-phase; M, M-phase of cell cycle. (**C**) Dot plot showing mRNA expression of membrane mucin genes in human colonic epithelial cell types. The color scheme shows row maximum and minimum representing the relative expression of each gene among all cell types in the intestinal segment. (**D**) Immunohistochemistry of MUC17 (green), Ezrin (magenta), and DNA (cyan) in histological sections of non-inflamed ileal biopsies from non-IBD control and CD patients, alongside semiquantitative analysis of MUC17 levels in the brush border. Each channel is also shown in grayscale. The yellow arrow points to the apical brush border. Scale bar 100 µm. n=16 for non-IBD, n=9 for CD. (**E**) Immunohistochemistry of MUC13 (green), Ezrin (magenta), and DNA (cyan) in histological sections of non-inflamed ileal biopsies from non-IBD control and CD patients, alongside semiquantitative analysis of MUC13 in the brush border. Each channel is also shown in grayscale. The yellow arrow points to the apical brush border. Scale bar 100 µm. n=16 for non-IBD, n=9 for CD. (**F**) Time-lapse glycocalyx permeability assay in non-inflamed ileal biopsies from non-IBD and CD patients, stained with CellMask, and incubated with *E. coli^GFP+^* tracked over the course of 6 min. Yellow arrows show points of contact between *E. coli^GFP+^* and CellMask. Dashed white lite marks the brush border. Magnifications are labeled with numbers. Scale bar 10 µm. The pseudo color scale indicates the time point during the tracking of individual bacterial cells. The bar graph depicts the frequency of *E. coli^GFP+^* attached to the brush border in each cohort. n=7 for non-IBD, n= 8 for CD. Data are means ± SD. Significance was determined by unpaired t-test (D-F). The estimation plot shows the difference between CD and non-IBD means with a 95% confidence interval.

Crohn’s disease (CD) is an IBD subtype characterized by patchy chronic inflammation involving dysregulated immune responses to the gut bacteria (*10*). Unlike UC that is limited to the colon, CD affects both the small and large intestines, with a third of patients displaying inflammation exclusively in the terminal ileum (*11*). Mutations in the intracellular peptidoglycan sensor *NOD2* and autophagy protein *ATG16L1* cause defects in the detection and degradation of intracellular bacteria and have been linked to susceptibility to CD (*12–14*). However, functional CD-associated defects upstream of intracellular innate immunity against bacteria have not been identified. Studies of ileal CD have reported transcriptional alterations associated with the enterocyte brush border (*15*). MUC17 is both transcriptionally and structurally associated with the brush border (*9*), but the role of MUC17 in epithelial defense and whether MUC17 expression and glycocalyx integrity are compromised in CD are not known.

Here, we analyzed non-inflamed ileal biopsies from CD patients and non-IBD controls together with a new genetic mouse model to define the function of MUC17 in the intestine. CD patients displayed decreased MUC17 levels and a glycocalyx that was more permeable to bacteria. In mice, intestine-specific deletion of the *Muc17* gene was dispensable for colonic inflammation whilst rendering the small intestine abnormally sensitive to infection by the attaching effacing bacterium *Citrobacter rodentium* (*C. rodentium*). *Muc17*^Δ*IEC*^ mice also suffered spontaneous deterioration of the epithelial cell barrier and displayed translocation of viable bacteria to extra-intestinal tissues. Lastly, we showed that Muc17 regulates the abundance of specific bacterial taxa in the small intestine, including bacteria enriched in CD patients. Thus, the *Muc17*^Δ*IEC*^ model reproduces fundamental manifestations of CD, suggesting that MUC17 is essential for protecting the small intestine against commensal and pathogenic bacteria.

### Decreased MUC17 levels and glycocalyx integrity in ileal Crohn’s disease

In humans, MUC17 is exclusively expressed in differentiated enterocytes of the small intestine (Fig. 1B-C). We analyzed the expression of MUC17 in histological sections from the non-inflamed terminal ileum of CD patients and a control group with non-IBD-related pathologies (table S1). MUC17 levels were significantly reduced in the enterocyte brush border of the CD compared to non-IBD ileum, which showed a typical MUC17 staining in the brush border marked by Ezrin (Fig. 1D). Membrane mucin MUC13 was not altered between the two cohorts (Fig. E). Next, we analyzed the ileal biopsies with an *ex vivo* time-lapse glycocalyx permeability assay (*9*), which measures the ability of the glycocalyx to separate GFP-expressing *Escherichia coli* (*E. coli^GFP+^*) from the brush border (see materials and methods). In line with decreased apical MUC17 in CD, the frequency of *E. coli^GFP+^* that approached the brush border was significantly higher in CD compared to non-IBD, indicating reduced glycocalyx barrier integrity in CD patients (Fig. 1F, fig. S1, Movie S1). Thus, the non-inflamed CD ileum exhibits an impaired glycocalyx barrier due to reduced MUC17 levels in the apical brush border.

### The *Muc17***^Δ^***^IEC^* mouse lacks a small intestinal glycocalyx

The loss of glycocalyx integrity in CD ileum suggests that the loss of apical MUC17 is an early defect resulting in increased bacterial contact with the brush border. To test this hypothesis, we generated a conditional *Muc17*^Δ*IEC*^ mouse by crossing *Muc17^fl/fl^* and *Vil1-Cre* mice expressing the Cre recombinase in intestinal epithelial cells (fig. S2A-C). Unlike human MUC17, murine Muc17 is expressed in both the small intestine and colon (Fig. 2A-B, fig. S3, fig. S4). The successful deletion of *Muc17* was thus confirmed in the duodenum, jejunum (Si5), ileum (Si8), proximal colon (PC), and distal colon (DC) (Fig. 2C, fig. S3, fig. S4).

**Fig. 2.**
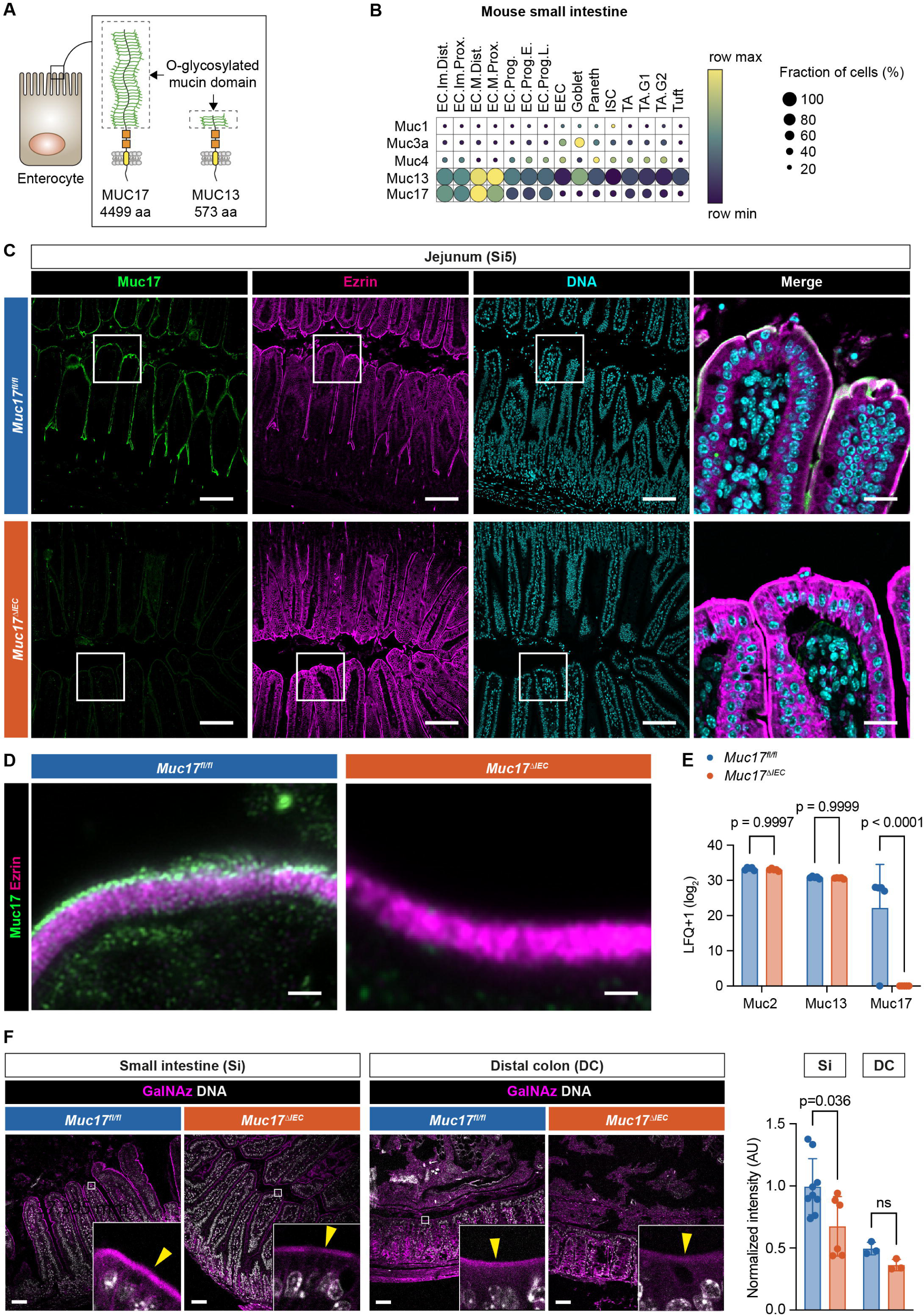
Epithelial-specific deletion of *Muc17* results in the loss of Muc17 expression in the mouse small intestine. (**A**) Schematic illustration of murine Muc17 and Muc13. Dashed boxes show the *O*-glycosylated mucin domain in each mucin. aa, amino acids. (**B**) Expression of membrane mucin transcripts in epithelial cell types of the mouse small intestine. The color scheme shows row maximum and minimum representing the relative expression of each gene among all cell types. EC, enterocyte; EEC, enteroendocrine cell; ISC, intestinal stem cell; TA, transit amplifying; Im, immature; M, mature; Prox, proximal; Dist, distal; Prog, progenitor; E, early; L, late; G1, G1-phase; G2, G2-phase of cell cycle. (**C**) Immunohistochemistry of Muc17 (green), Ezrin (magenta), and DNA (cyan) in histological sections of the jejunum (Si5) from *Muc17^fl/fl^* and *Muc17*^Δ*IEC*^ mice. Scale bar 50 µm. Scale bar in insets 10 µm. (**D**) Airyscan high-resolution microscopy of Muc17 (green) and Ezrin (magenta) in the small intestinal brush border of *Muc17^fl/fl^*and *Muc17*^Δ*IEC*^ mice. Scale bar 1 µm. (**E**) Bar graph representing the abundance of detected mucins in the proteome of small intestinal epithelial cells from *Muc17^fl/fl^* and *Muc17*^Δ*IEC*^ mice. n= 5 for each group. (**F**) Representative confocal micrographs of small intestine (Si) and distal colon (DC) from *Muc17^fl/fl^* and *Muc17*^Δ*IEC*^ mice, stained for bioorthogonally labeled *O*-glycans (GalNAz, magenta) and DNA (white). Scale bar 50 µm. Bar graphs represent the quantification of GalNAz intensity in the apical brush border of the small intestine (Si) and DC in each group. Si5: n=6-9 per group. DC: n=3 per group. Significance was determined by two-way ANOVA followed by Šidák correction (D) and Mann-Whitney test (E).

High-resolution microscopy of small intestinal enterocytes revealed that MUC17 was lacking from the tips of microvilli in *Muc17*^Δ*IEC*^ mice. (Fig. 2D). *Muc17* deletion was also demonstrated in microvillus-derived luminal vesicles isolated from the small intestinal (*16, 17*) (fig. S2D). To evaluate the impact of *Muc17* deletion on other mucins, we performed proteomic analysis of isolated small intestinal epithelial cells from *Muc17^fl/fl^* and *Muc17*^Δ*IEC*^ cohoused littermates. Deletion of *Muc17* did not result in a compensatory change in the abundance of Muc13 or the gel-forming Muc2 (Fig. 2E, table S2). Moreover, *Muc17*^Δ*IEC*^ mice did not exhibit alterations in tissue morphology of the small intestine or distal colon (fig. S2E). Notably, the deletion of *Muc17* did not cause abnormalities in brush border morphology or cellular polarity, evaluated by the brush border markers Ezrin and Cdhr5, and basolateral EpCAM (fig. S2F-G). Mucin *O*-glycans convert the glycocalyx into a size-selective diffusion barrier and shield against bacterial degradation (*1, 9*). To determine if the loss of Muc17 impacted the density of glycocalyx *O*-glycans, we employed *in vivo* bioorthogonal labeling using GalNAz, the azide derivative of N-Acetylgalactosamine (GalNAc), the first glycan added to serines or threonines during mucin-type *O*-glycosylation (*18*). GalNAz intensity was significantly reduced in the small intestinal brush border of *Muc17*^Δ*IEC*^ mice compared to controls but remained unchanged in the colonic brush border (Fig. 2F). The *O*-glycan deficit of the small intestinal glycocalyx was further supported by decreased staining with Aleuria aurantia lectin (AAL) and wheat germ agglutinin (WGA), which bind fucose and N-acetylglucosamine (fig. S2H). Thus, under baseline conditions *Muc17* deletion results in an explicit loss of the glycocalyx without impacting the overall tissue and cell morphology of the small intestine.

### Deletion of *Muc17* causes abnormal small intestinal susceptibility to *Citrobacter Rodentium* infection

Since we did not observe any baseline phenotype in *Muc17*^Δ*IEC*^ mice, cohoused *Muc17^fl/fl^* and *Muc17*^Δ*IEC*^ littermates were infected with *C. rodentium*, a natural murine pathogen that normally causes a self-limiting infection in the distal colon (*19*). Infection kinetics in each group were monitored over a period of 21 days post-infection (dpi) (Fig. 3A). Regardless of genotype, fecal *C. rodentium* counts peaked at 7-9 dpi and declined below the limit of detection (LOD) by 17 dpi (Fig. 3B). Both groups exhibited a slight but comparable weight loss during the initial infection phase (1-3 dpi) (Fig. 3C). Next, we quantified *C. rodentium* colonization in the luminal and mucosal compartments of the jejunum (Si5), ileum (Si8) and distal colon (DC) at 3 days post-infection. As expected, *Muc17^fl/fl^* mice exhibited a robust colonization in DC (Fig. 3D-E). In *Muc17*^Δ*IEC*^ Si5, however, we observed three orders of magnitude higher *C. rodentium* counts compared to the control group (Fig. 3D-E). The abnormal *C. rodentium* infection of Si5 was not limited to the luminal compartment but was also detected in the mucosa, indicating that the small intestinal epithelium was more susceptible to *C. rodentium* infection in the absence of Muc17 and a glycocalyx. Greater pathogen counts were also observed in the mucosa of *Muc17*^Δ*IEC*^ DC compared to the infected controls (Fig. 3E). There was also a trend towards increased translocation of *C. rodentium* to the peripheral tissues in *Muc17*^Δ*IEC*^ mice at 7 dpi (Fig. 3F). Notably, across all tissues and time points post-infection, a larger fraction of *Muc17*^Δ*IEC*^ mice carried a detectible pathogen burden compared to *Muc17^fl/fl^* mice (fig. S5). At 7 days post-infection, 91±6 % of all *Muc17*^Δ*IEC*^ mice exhibited a small intestinal infection compared to only 54±13 % of the *Muc17^fl/fl^* mice. A similar analysis of colonic pathogen burden did not reveal any significant difference between the groups. Accordingly, the relative risk of *C. rodentium* infection in *Muc17*^Δ*IEC*^ mice peaked in Si5 and decreased towards the DC, suggesting that Muc17 primarily serves as a small intestinal defense mechanism during bacterial infection (Fig. 3G). *Muc17* deletion also resulted in a higher relative risk of pathogen translocation to the peripheral tissues (Fig. 3H). Histological analysis 3 days post-infection revealed numerous *C. rodentium^GFP+^* cells in contact with the epithelium at villus apexes, intervillar regions, and the crypt compartment of *Muc17*^Δ*IEC*^ Si5, whereas *C. rodentium^GFP+^* was not detected in *Muc17^fl/fl^* (Fig. 3I). Assessment of the distal colon showed *C. rodentium^GFP+^* at the surface epithelium and in the crypts of *Muc17*^Δ*IEC*^ mice (Fig. 3J-K). Collectively, the absence of Muc17 caused an abnormal small intestinal susceptibility to *C. rodentium*, indicating that Muc17 is essential for protecting the small intestinal epithelium against pathogenic bacteria.

**Fig. 3.**
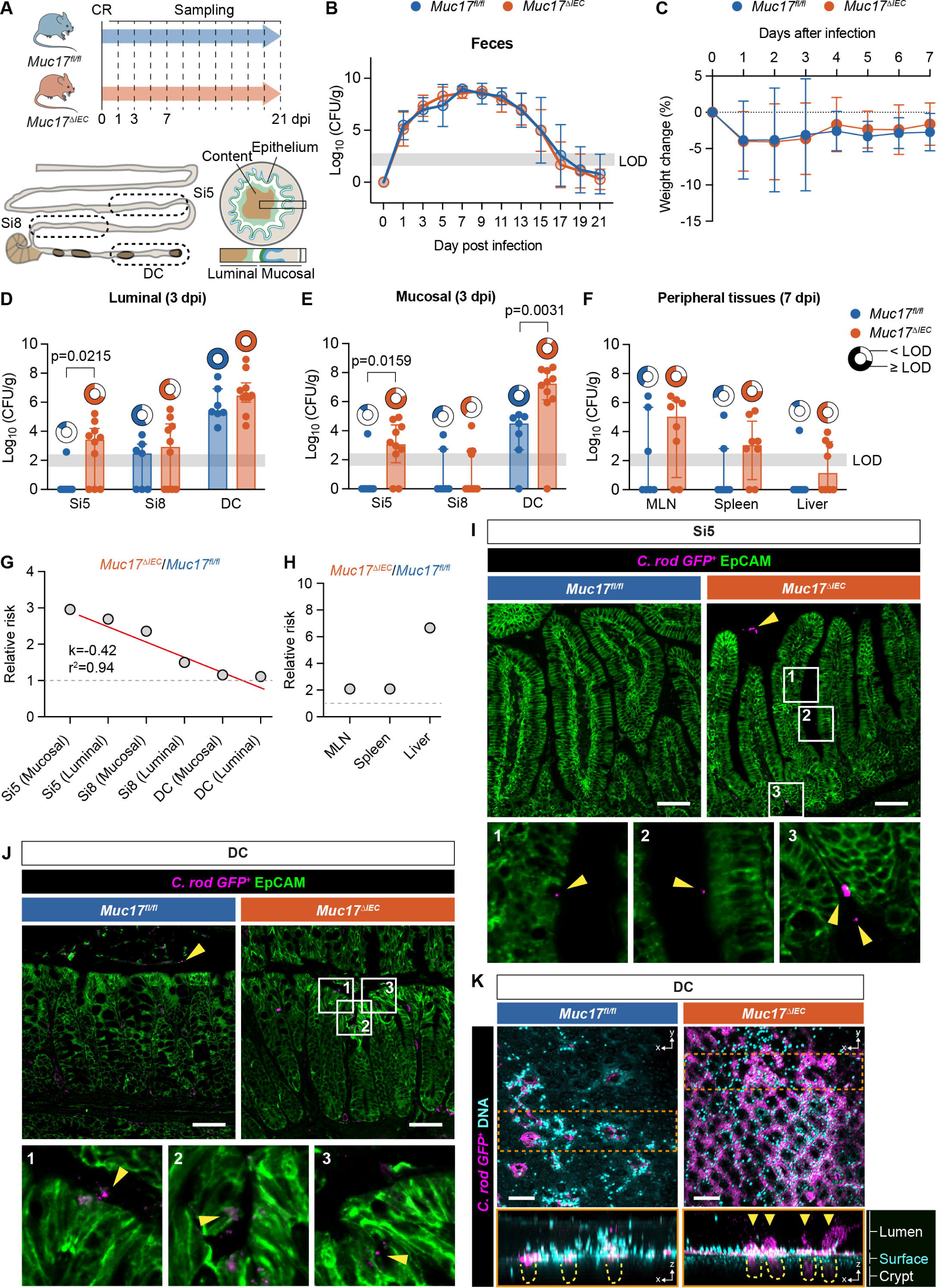
Deletion of Muc17 results in abnormal small intestinal susceptibility to *C. rodentium* infection. (**A**) Schematic illustration of the *C. rodentium* infection protocol and sampling time points. Si5, jejunum; Si8, ileum; DC, distal colon. (**B**) Colony forming units (CFU) of *C. rodentium* in fecal samples collected from *Muc17^fl/fl^* and *Muc17*^Δ*IEC*^ mice at various days post-infection (dpi). (**C**) Weight loss of infected *Muc17^fl/fl^* and *Muc17*^Δ*IEC*^ mice, depicted as percent of the initial weight at the start of the challenge (time point 0 dpi). (**D**) CFU of *C. rodentium* in the luminal compartments of the jejunum (Si5), ileum (Si8), and distal colon (DC) at 3 dpi. (**E**) CFU of *C. rodentium* the mucosal compartments of the jejunum (Si5), ileum (Si8), and distal colon (DC) at 3 dpi. (**F**) CFU of *C. rodentium* in the mesenteric lymph nodes (MLN), Spleen, and Liver at 7 dpi. The circle charts represent the proportion of mice with each genotype carrying *C. rodentium* CFU above the limit of detection (LOD) in each intestinal segment or tissue site. (**G**) Relative risk of carrying *C. rodentium* CFU above LOD in Si5, Si8, and DC when *Muc17*^Δ*IEC*^ and *Muc17^fl/fl^* mice are challenged with *C. rodentium*. (**H**) Relative risk of carrying *C. rodentium* CFU above LOD in MLN, Spleen, and liver when *Muc17*^Δ*IEC*^ and *Muc17^fl/fl^* mice are infected with *C. rodentium*. n=6-10 per group. (**I**) Immunohistochemistry of *C. rodentium^GFP+^* (magenta) in relation to the epithelium (EpCAM, green) in Si5 of *Muc17^fl/fl^* and *Muc17*^Δ*IEC*^ mice 3 days post-infection. Yellow arrows point to bacterial cells. Scale bar 50 µm. (**J**) Immunohistochemistry of *C. rodentium^GFP+^* (magenta) in relation to the epithelium (EpCAM, green) in DC of *Muc17^fl/fl^* and *Muc17*^Δ*IEC*^ mice 3 days post-infection. Yellow arrows point to bacterial cells. Scale bar 50 µm. (**K**) Visualization of *C. rodentium^GFP+^* (magenta) in relation to the surface epithelium (DNA, cyan) in explants of *Muc17^fl/fl^* and *Muc17*^Δ*IEC*^ DC 3 days post-infection. Upper panels show a top view (xy plane), and lower panels show an extended orthogonal view (xz plane) of the boxed region (orange). Penetration of *C. rodentium^GFP+^* (magenta) into the colonic crypts (yellow dashed lines) is highlighted (yellow arrows). Scale bar 100 µm. Significance was determined by Mann-Whitney test (D-F).

### *Muc17***^Δ^***^IEC^* mice display spontaneous loss of epithelial homeostasis

Given that Muc17 protected the small intestine against pathogenic infection, we reasoned that *Muc17* deletion might also allow commensal bacteria to make direct contact with enterocytes and thereby disrupt epithelial homeostasis. Thus, we turned our attention to unchallenged 6-8-week-old cohoused *Muc17^fl/fl^* and *Muc17*^Δ*IEC*^ littermates maintained under standard pathogen-free (SPF) conditions. Histological analysis of Si5 did not reveal any genotype-dependent differences in crypt number, villus length, or goblet cell frequency (Fig. 4A, fig. S2E). Next, Si5 explants were flushed, and fixed to preserve the mucosal compartment, and the distribution of bacteria in relation to the brush border was assessed using confocal microscopy and automatic image segmentation. Compared to *Muc17^fl/fl^*, significantly higher numbers of bacteria were directly bound to the brush border of *Muc17*^Δ*IEC*^ mice (Fig. 4B). Findings were supported by fluorescence *in situ* hybridization (FISH) with a universal eubacterial probe, showing bacteria in direct contact with the *Muc17*^Δ*IEC*^ brush border (Fig. 4C). Uncontrolled bacterial interactions with the epithelium can trigger increased epithelial proliferation and cell death (*20, 21*). Correspondingly, we observed a significant expansion of proliferative mKi67^+^ epithelial cells within the crypt compartment of *Muc17*^Δ*IEC*^ Si5 (Fig. 4D). Free nuclear DNA 3′-OH termini generated by single-strand breaks are hallmarks of apoptosis. Compared to controls, *Muc17*^Δ*IEC*^ Si5 displayed a higher number of TUNEL-positive apoptotic cells at villus apexes (Fig. 4E). Next, we used mass spectrometry to evaluate the impact of *Muc17* deletion on epithelial homeostasis. Label-free protein quantification in epithelial cells from 5-, 8- and 36-weeks-old *Muc17^fl/fl^* and *Muc17*^Δ*IEC*^ mice revealed altered expression in 0.25-1.00% of the proteome, mainly affecting brush border-associated proteins (Fig. 4F, table S2). Specifically, epithelial cells in 8-week-old *Muc17*^Δ*IEC*^ mice overexpressed Rho guanine nucleotide exchange factor 26 (Arhgef26), which participates in enterocyte membrane ruffling during bacterial invasion (*22*) (Fig. 4F, fig. S6). In line with the persistent bacterial challenges of the epithelium, we also detected higher abundance of the antimicrobial proteins Lyz1 and Reg3b in 36-week-old *Muc17*^Δ*IEC*^ mice (Fig. 4F, fig. S6).

**Fig. 4.**
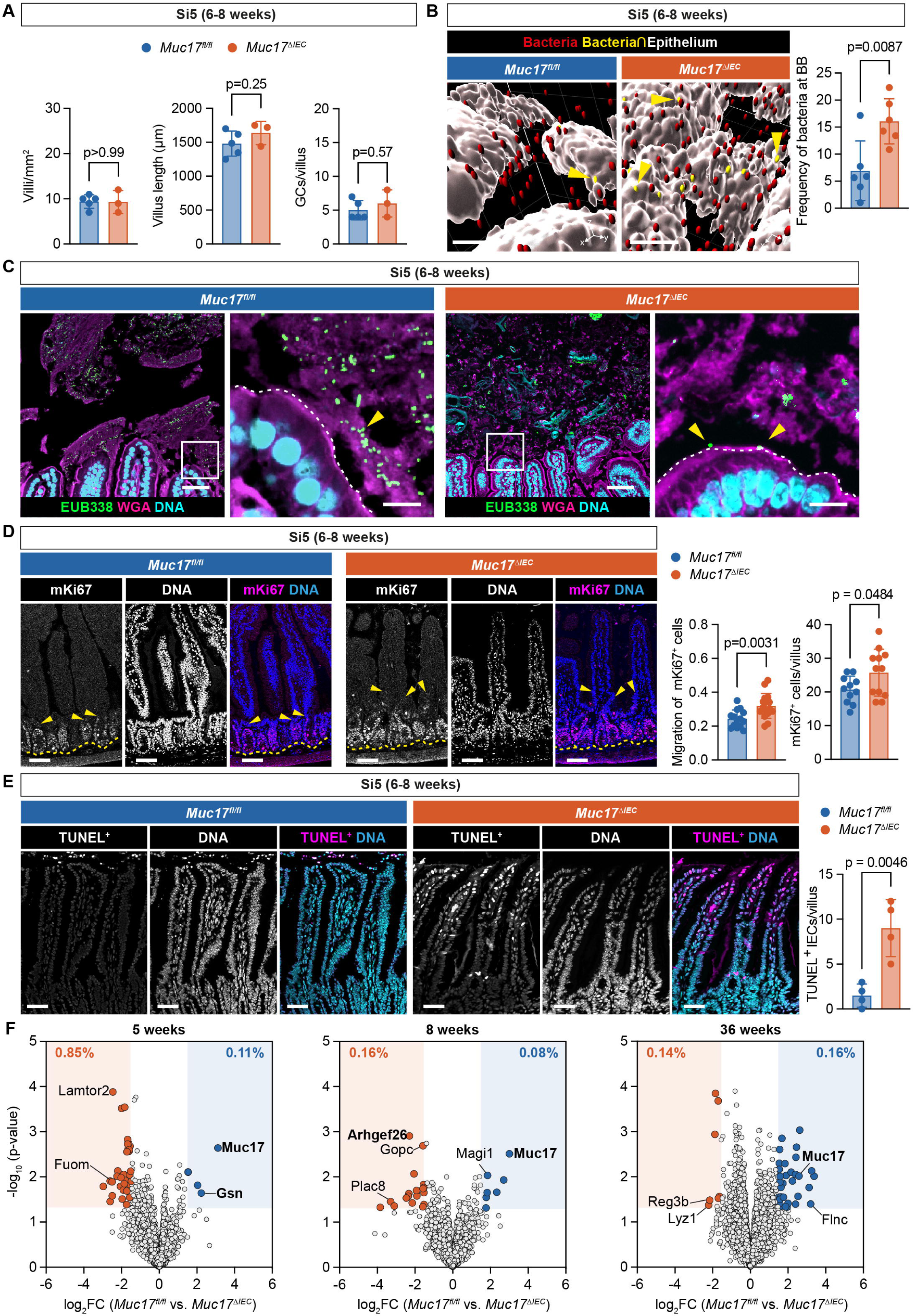
Epithelial barrier dysfunction in the small intestine of *Muc17***^Δ^***^IEC^* mice housed under baseline SFP conditions. (**A**) Quantification of the number of villi per mm^2^ of histological section, villus length, and goblet cells (GCs) per villus in *Muc17^fl/fl^* and *Muc17*^Δ*IEC*^ jejunum (Si5). n=5 for *Muc17^fl/fl^*, n=3 for *Muc17*^Δ*IEC*^. (**B**) Representative visualization of commensal bacteria (red) at the Si5 epithelium (white). Bacteria in direct contact with the epithelium are shown in yellow. Scale bar 100 µm. The quantification shows frequency of bacteria attached to the brush border in each cohort. N=6 for *Muc17^fl/fl^*, n=6 for *Muc17*^Δ*IEC*^. (**C**) Confocal micrographs of *Muc17^fl/fl^*and *Muc17*^Δ*IEC*^ Si5 stained for bacteria (EUB338, green), epithelium and mucus (WGA, magenta), and DNA (cyan). Yellow arrows point to bacteria. Scale bar 25 µm. Scale bar in insets 10 µm. (**D**) Immunohistochemistry of mKi67 (magenta) and DNA (blue) in *Muc17^fl/fl^* and *Muc17*^Δ*IEC*^ Si5. Each channel is also shown in grayscale. Yellow arrows point to mKi67^+^ cells with maximum migration along the villus-crypt axis. The dashed yellow line depicts the crypt bottom. Scale bar 50 µm. Quantification of the migration of mKi67^+^ cells along the crypt-villus axis and absolute numbers of mKi67^+^ per villus in *Muc17^fl/fl^* and *Muc17*^Δ*IEC*^ Si5. n=3-6 villi per mouse, 3 mice per group. (**E**) Immunohistochemistry of TUNEL^+^ nuclei (magenta) and DNA (cyan) in *Muc17^fl/fl^* and *Muc17*^Δ*IEC*^ Si5. Each channel is also shown in grayscale. Scale bar 50 µm. Quantification of TUNEL^+^ cells per villus in *Muc17^fl/fl^* and *Muc17*^Δ*IEC*^ mice. n=4 for *Muc17^fl/fl^*, n=4 for *Muc17*^Δ*IEC*^. **(F)** Volcano plots showing the fold change of protein expression in epithelial cells of the jejunum (Si5) from 5-, 8-, and 36-week-old *Muc17^fl/fl^* compared to *Muc17*^Δ*IEC*^ mice. Differentially expressed proteins (fold change ≥ 2, p-value < 0.05) are highlighted with filled circles. Specific proteins are labeled with their protein names. n=5 for each genotype in each age group. Significance was determined by Mann-Whitney test (A) and unpaired t-test (B, D, E).

### Muc17 is dispensable for protection against chemically induced colitis

To define the protective role of Muc17 in the murine distal colon, cohoused *Muc17^fl/fl^* and *Muc17*^Δ*IEC*^ littermates were subjected to 7-day *ad libitum* administration of dextran sodium sulfate (DSS), a chemically induced model of human ulcerative colitis (fig. S7A). *Muc17*^Δ*IEC*^ mice displayed a reduction in weight compared to wild-type mice, but only at the end of the intervention (day 6). (fig. S7B). While there was a significant shortening of the colon in *Muc17*^Δ*IEC*^ mice after 7 days (fig. S7C), only a moderately increased susceptibility to DSS was measured by stool consistency, fecal blood score, overall disease activity index, and survival (fig. S7D-G). Our observations were supported by histological analysis of Si5 and DC, which corroborated that *Muc17^fl/fl^* and *Muc17*^Δ*IEC*^ mice responded similarly to DSS-induced colitis (fig. S7H-I). The late onset of moderate inflammation in the absence of Muc17 contrasted acute induction of severe colitis in *C3GnT^−/−^*, *Vamp8^−/−^*, and *Tgm3^−/−^*mice with defects in Muc2 glycosylation, secretion, and crosslinking (*23–25*). Thus, we concluded that Muc17 does not play a critical protective role in the distal colon.

Further assessment of *Muc17*^Δ*IEC*^ mice under baseline conditions did not show any deviations in the number of crypts, crypt length, or goblet cell frequency in the distal colon (fig. S8A, fig. S2E). Next, we quantified the barrier properties of the inner mucus layer (IML) separating bacteria from the epithelium (*26*). FISH showed a defined separation of bacteria from the epithelium in 6-8-week-old *Muc17^fl/fl^* and *Muc17*^Δ*IEC*^ mice (fig. S8B*). Ex vivo* mucus penetrability analysis in viable explants from 6–8- or 36-40 week-old mice, evaluating the penetration of microbeads through the IML, showed that both *Muc17*^Δ*IEC*^ and *Muc17^fl/fl^* mice maintain an impenetrable IML typically observed in wild-type mice (*3*) (fig. S8C-D). *Muc17*^Δ*IEC*^ mice did however display elevated occurrence of shed cells (fig. S8D) and increased epithelial cell proliferation measured by mKi67^+^ (fig. S8E). Together, our findings confirm that Muc17 is not required for the protection of the distal colon against commensal bacteria.

### *Muc17* deletion alters the small intestinal microbiota and elicits extra-intestinal translocation of bacteria

*Muc17* deletion resulted in susceptibility to *C. rodentium* and increased contact between commensal bacteria and small intestinal enterocytes, but the precise effect of Muc17 on the regiospecific composition of the microbiota is not understood. We thus analyzed bacterial genomic DNA from the luminal compartment of the small intestine and colon of cohoused *Muc17^fl/fl^* and *Muc17*^Δ*IEC*^ littermates by 16S rRNA gene sequencing. While there was no significant difference in species richness and evenness (alpha diversity) between the two groups (Fig. 5A and 5D), the beta diversity based on Bray-Curtis distances revealed distinct patterns in the microbiota of *Muc17^fl/fl^* and *Muc17*^Δ*IEC*^ mice (Fig. 5B and 5E). The small intestinal community of *Muc17*^Δ*IEC*^ mice was characterized by a higher abundance of the classes of *Actinobacteria*, *Bacteroidia*, *Coriobacteria*, and *gammaproteobacteria* (Fig. 5C), while the colonic microbiota shifted towards *Clostridia*, *Coriobacteria*, and *Desulfovibrionia* (Fig. 5F). Linear discriminant analysis effect size (LEfSe) analysis showed a depletion of *Firmicutes*, *Bacilli* and the genus *Dubosiella* belonging to the *Erysipelotrichales* from the *Muc17*^Δ*IEC*^ small intestine (Fig. 5G, fig. S9A). The small intestinal alterations in composition were largely reproduced in the *Muc17^fl/fl^* colon, with the enrichment of *Erysipelotrichales* and *Dubosiella* alongside *Lactobacillus* (Fig. 5G, fig. S9B). In addition, *Coriobacteriales*, *Lachnoclostridium*, and *Marvinbryantia* were enriched in the *Muc17*^Δ*IEC*^ colon (Fig. 5G, fig. S9B). Combined, the affected genera comprised an average of 15%-20% of the detected operational taxonomic units. Since the microbiota composition varies along the intestine (*27*), we assessed the regional distribution of the differentially enriched bacterial taxa. *Bacilli*, *Coriobacteriia*, and *Erysipelotrichales,* had a primarily small intestinal niche, with *Erysipelotrichales* dominating the jejunum (Si5) (Fig. 5G), thus suggesting a transmission of compositional changes from the small intestine to colon. Collectively, *Muc17* deletion induced a substantial shift in the abundance of specific bacteria that colonize the small intestine (fig. S9C), suggesting that Muc17 regulates the regiospecific selection of commensal bacteria.

**Fig. 5.**
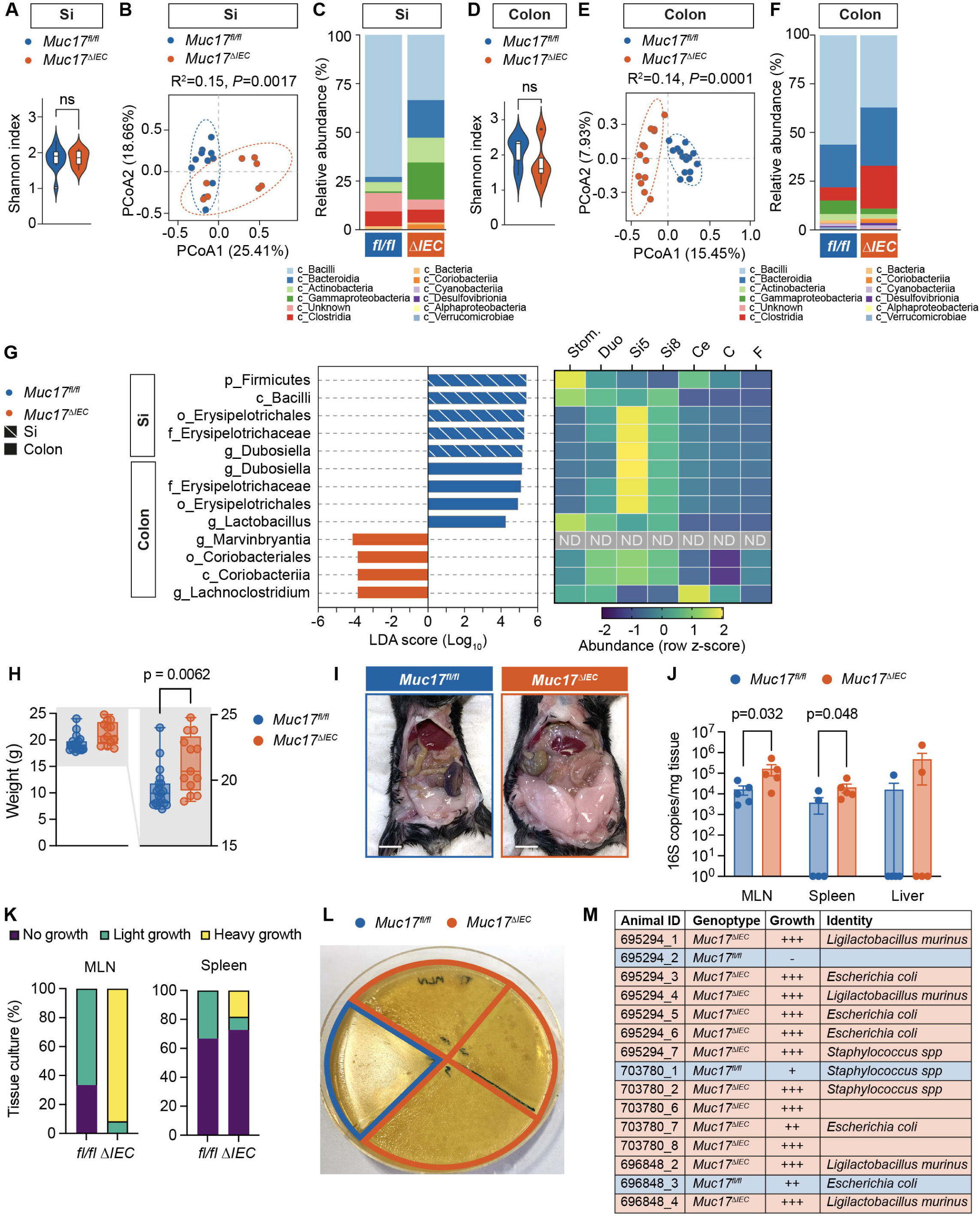
Deletion of Muc17 results in alterations in luminal microbiota composition and translocation of commensal bacteria to peripheral tissues. (**A**) α-diversity (Shannon index) of luminal bacterial communities in the small intestine (Si) of *Muc17^fl/fl^* (n=11) and *Muc17*^Δ*IEC*^ (n=8) mice. (**B**) Principal coordinate analysis of β-diversity (Bray-Curtis dissimilarity) of luminal bacterial communities of *Muc17^fl/fl^* and *Muc17*^Δ*IEC*^ Si. (**C**) Relative frequency of major classes found in *Muc17^fl/fl^* and *Muc17*^Δ*IEC*^ Si. (**D**) α-diversity (Shannon index) of luminal bacterial communities in the colon of *Muc17^fl/fl^*(n=15) and *Muc17*^Δ*IEC*^ (n=13) mice. (**E**) Principal coordinate analysis of β-diversity (Bray-Curtis dissimilarity) of luminal bacterial communities in *Muc17^fl/fl^* and *Muc17*^Δ*IEC*^ colon. (**F**) Relative frequency of major classes found in *Muc17^fl/fl^* and *Muc17*^Δ*IEC*^ colon. (**G**) Linear discriminant analysis (LDA) effect size identification of small intestinal and colonic taxa that differentiate between *Muc17^fl/fl^* (blue) and *Muc17*^Δ*IEC*^ (red) mice. The heat map shows the row z-score representing the relative abundance of the selected taxa in different gastrointestinal segments. Stom, stomach; Duo, duodenum; Si5, jejunum; Si8, ileum; Ce, cecum; C, colon; F, feces; ND, not detected. (**H**) Total weight of cohoused *Muc17^fl/fl^* (n=19) and *Muc17*^Δ*IEC*^ (n=13) littermate mice. Box plots show the median and minimum to maximum values for each group. (**I**) Representative image of the opened abdomen of 36-40-week-old *Muc17^fl/fl^* and *Muc17*^Δ*IEC*^ mice. Scale bar 1 cm. (**J**) Quantification of bacterial 16S copy number in mesenteric lymph nodes (MLN), spleen, and Liver tissue by qPCR. n= 5 for each group. (**K**) Quantification of bacterial growth in tissue homogenates of MLN and spleen on BHIS agar plates under anaerobic conditions. (**L**) Representative image representing bacterial growth in tissue homogenates of MLN on BHIS agar under anaerobic conditions. (**M**) Identification of isolated bacterial colonies identified by 16S rDNA sequencing. Significance was determined by unpaired t-test (A) and Mann-Whitney test (C).

In ileal Crohn’s disease, displacement of mucosa-associated bacteria from the site of inflammation to mesenteric adipose tissue induces the expansion of encapsulating “creeping fat” (*28*). Bacterially induced enlargement of visceral adipose tissue also occurs in mice that fail to control microbial translocation (*29*). Under baseline SPF conditions, *Muc17*^Δ*IEC*^ mice displayed a higher total body mass (Fig. 5H) and larger abdominal fat pads compared to age-matched *Muc17^fl/fl^* cohoused littermates (Fig. 5I). Subsequently, we asked if *Muc17*^Δ*IEC*^ mice displayed translocation of commensal bacteria to extra-intestinal tissues. Absolute 16S rRNA gene quantification uncovered significantly higher counts in the MLNs and spleen of *Muc17*^Δ*IEC*^ mice compared to wild-types (Fig. 5J). Bacterial quantification was complemented with anaerobic culturing of viable bacteria from the affected tissues. Heavy bacterial growth was observed in 92% of *Muc17*^Δ*IEC*^ MLNs, with the remaining 8% exhibiting light growth (Fig. 5K-L). In *Muc17^fl/fl^* mice, 33% of MLNs were clear of bacteria and 67% exhibited only light growth. We also observed a higher number of *Muc17*^Δ*IEC*^ spleens with heavy bacterial growth compared to control tissues (Fig. 5K). Sequencing of the 16S gene identified the translocated bacteria as *Escherichia coli*, *Staphylococcus* spp., and *Ligilactobacillus murinus* (Fig. 5M), the latter described as a translocating strain (*30*). Thus, we concluded that viable commensal bacteria penetrate the intestinal epithelial cell barrier in the absence of Muc17, highlighting its key role in small intestinal barrier function.

## Discussion

It is increasingly understood that the defensive properties of gel-forming mucins are important in protection against the development of human IBD (*4*), but the role of membrane mucins still remains elusive. In this study, we used a conditional mouse model to identify Muc17 as essential for the regiospecific protection of the small intestine against commensal and pathogenic bacteria. Significantly, a reduction in MUC17 levels in the enterocyte brush border leading to impaired glycocalyx protection against bacteria, was observed already in the non-inflamed ileum of CD patients. Our findings suggests that membrane mucin dysfunction precedes inflammation in CD.

The mechanisms underlying CD include genetic factors, such as *NOD2* mutations that impair the sensing and removal of intracellular bacteria (*10*). These defects in turn trigger immune responses that evoke chronic inflammation. While there are no known disease-associated mutations in the *MUC17* gene, studies have reported an altered glycocalyx ultrastructure in ileal CD (*15*). However, the implications of a compromised glycocalyx and the loss of its major component MUC17 in CD have not been addressed. In this study, we identified decreased MUC17 levels and a weakened glycocalyx in ileal CD to allow for increased bacteria-epithelium interactions. The results find mechanistic support in our earlier studies of murine small intestinal explants demonstrating that Muc17 forms the enterocyte glycocalyx (*9*). We also showed that the formation of the glycocalyx by Muc17 is induced by IL-22, a critical regulator of antibacterial defenses, including antimicrobial peptides (Reg3b and Reg3g) and mucin fucosylation via *FUT2* (*31*). Notably, many IBD risk genes, such as *IL23R*, *JAK1/2*, *TYK2*, and *STAT3*, participate in IL-22 signaling (*31*) and the frequency of IL-22-producing type 3 innate lymphoid cells is lower in CD patients (*32*).

Human MUC17 is confined to small intestinal enterocytes, while murine MUC17 is expressed in both the small and large intestines. We disentangled the segment-specific function using two interventions with distinct mechanisms of action. DSS disrupts the mucus layer separating gut bacteria from the epithelium (*33*). Accordingly, DSS-induced colitis is rapid and severe in mouse models with mucus defects (*23–25*). Since glycocalyx-deficient *Muc17*^Δ*IEC*^ mice still produced a functional colonic mucus barrier, they were equally sensitive to colitis as wild-type controls. In an alternative intervention, we used *C. rodentium*, which shares virulence factors with enteropathogenic *Escherichia coli* (EPEC), a major human pathogen that targets the small intestine. While *C. rodentium* is typically used to study pathogen-host interactions in the mouse distal colon (*34*), it surprisingly colonized the jejunum, including its mucosa, in Muc17-deficient mice. This finding highlights the crucial role of MUC17 in defending the small intestine against bacterial infections, even in the presence of a functioning mucus barrier. Interestingly, the ileum of Muc17-deficient mice was more resistant to colonization by *C. rodentium* than the jejunum. This difference can be explained by the lower expression of antimicrobial peptides in the mouse jejunum compared to the ileum, making it easier for the bacteria to colonize the jejunum (*35*). Given the link between CD and reduced antimicrobial peptide levels (*36, 37*), the murine jejunum emerges as a relevant model for ileal CD in humans.

Defects in intestinal defenses frequently result in the translocation of gut bacteria to peripheral tissues (*26, 29*). Bacterial translocation has also recently been linked the expansion of “creeping fat” in CD (*28*). Under baseline conditions, *Muc17*^Δ*IEC*^ mice exhibited a spontaneous deterioration of epithelial homeostasis in the small intestine, involving elevated levels of host proteins participating in bacterial invasion and antimicrobial defenses. That *Muc17*^Δ*IEC*^ mice displayed greater body mass, enlargement of abdominal fat, and viable commensal bacteria in peripheral tissues further supports the importance of the glycocalyx-forming MUC17 in limiting pathology related to CD.

Gel-forming mucins not only serve as a physical barrier against bacteria but also provide a habitat for bacteria that utilize mucins as a carbon source (*35, 38*). However, such a role for membrane mucins has not been described. Our work shows that Muc17 regulates gut microbiota composition and signifies that Muc17-dependent changes in the bacterial community propagate from the small intestine, where Muc17 exerts its function, distally towards the colon. Importantly, increased abundance of *Coriobacteriia* and *Lachnoclostridium*, enriched in *Muc17*^Δ*IEC*^ mice, has also been reported in CD patients (*39–41*). Thus, our study suggests that MUC17 fulfills two purposes; forming a glycocalyx that blocks bacteria and regulating the selection of bacteria by the host. We postulate that the latter function depends on the apical localization of MUC17 on microvilli, which release luminal vesicles that bind and limit bacterial growth in the lumen (*16*). The absence of Muc17 from microvillus-derived luminal vesicles could alter vesicle interactions with bacteria, thereby impacting the microbial community.

In conclusion, we uncovered a role for the MUC17-based glycocalyx as an important defense system of the small intestine. Our findings shed light on the regiospecific role of membrane mucins as innate defense components that create a critical interface between the host epithelium and the gut microbiota. The disruption of this system in the non-inflamed ileum of CD patients suggests that defective MUC17 biosynthesis or trafficking is an early epithelial defect that precedes inflammation. Given the limited number of mouse models for small intestinal inflammation, the *Muc17*^Δ*IEC*^ model provides opportunities for understanding the molecular mechanisms underlying epithelial cell dysfunction in Crohn’s disease.

## Acknowledgements

The authors acknowledge support from the National Genomics Infrastructure in Genomics Production Stockholm funded by Science for Life Laboratory, the Knut and Alice Wallenberg Foundation and the Swedish Research Council (2017.2008), and SNIC/Uppsala Multidisciplinary Center for Advanced Computational Science for assistance with massively parallel sequencing and access to the UPPMAX computational infrastructure. We also thank the Bioinformatics and Data Centre at the Sahlgrenska Academy and Clinical Genomics Gothenburg at SciLifeLab for bioinformatics support. The authors also thank all the helpful colonoscopists at the Endoscopy Unit of the Sahlgrenska University Hospital.

## Funding

Swedish Society for Medical Research (Svenska Sällskapet för Medicinsk Forskning) S17-0005 (TP)

National Institutes of Health 5U01AI095542-08-WU-19-95, 5U01AI095542-09-WU-20-77 (TP)

Wenner-Gren Stiftelserna FT2017-0002 (TP) Jeansson Foundations JS2017-0003 (TP)

Åke Wiberg Foundation M17-0062, M21-0022 (TP) Biocodex Microbiota Foundation (TP)

Stiftelsen Clas Groschinskys Minnesfond M2254 (TP) Sahlgrenska Academy International Starting Grant E2015/521 (TP)

Grants under the ALF agreement 236501 between the Swedish Government and the county councils, ALFGBG-991311, ALFGBG-440741 (TP)

Wenner-Gren Foundations UPD2018-0065, WUP2017-0005 (ELH)

Wilhelm och Martina Lundgrens Vetenskapsfond 2020-3647, 2021-3880, 2022-4051 (ELH) Sahlgrenska Academy Core Facilities Elisabeth “Bollan” Lindén Stipend (ELH)

B.A.V. is the Children with Intestinal and Liver Disorders Foundation Chair in Pediatric Gastroenterology

## Author contributions

Conceptualization: ELH, TP

Methodology: ELH, SJ, LA, BD, TP

Investigation: ELH, SJ, FCvdK, MS, TP

Visualization: ELH, TP

Supervision: TP Materials: ST, BAV, TP

Writing—original draft: ELH, TP

Writing—review & editing: ELH, SJ, FCvdK, BD, LA BAV, TP

## Data and materials availability

All data are available in the main text or the supplementary material. Primary data for all the described experiments are reported in table S4. The mass spectrometry proteomics data have been deposited to the ProteomeXchange Consortium (http://proteomecentral.proteomexchange.org) via the PRIDE partner repository with the dataset identifier PXDXXXX.

