## Supplementary materials for "MUC17 is an essential small intestinal glycocalyx component that is disrupted in Crohn’s disease"

Elena Layunta *et al.*

**This PDF file includes:**

Material and Methods

Figs. S1 to S9

Tables S1 to S4

Movie S1

**Materials and Methods**

**Experimental Design**

We aimed to identify the molecular mechanisms by which compromised MUC17 and glycocalyx function impact disease development in ileal CD. To this end, non-inflamed ileal biopsies from CD and non-IBD patients were analyzed for MUC17 levels and glycocalyx barrier integrity. Mechanistic and functional investigations were performed using a novel preclinical *Muc17^∆IEC^* mouse strain, which was subjected to two distinct challenge models as well as analyzed longitudinally under baseline SFP conditions. Data collection and analysis were performed in a blinded manner whenever possible. For patient biopsies, sample size was not determined by power calculation but estimated based on our experience and previous studies. Sample sizes and statistical tests are described in the figure legends.

**Human subjects**

Patients (≥18 years) with Crohn’s disease (CD) and individuals without suspected IBD (non-IBD), referred to Sahlgrenska University Hospital (Gothenburg, Sweden), were eligible for inclusion in the study. Informed consent was obtained from all patients in accordance with the respective protocol described in the ethical permit 2020-03196. Clinical information and metadata for enrolled subjects are provided in table S1. Patients with macroscopic or microscopic evidence of intestinal pathology other than CD were excluded. Biopsies were obtained during ileocolonoscopy, using biopsy forceps routinely used in standard of care. Three biopsies were obtained from the non-inflamed terminal ileum of each subject and immediately placed into the appropriate vials; ice-cold oxygenated Krebs transport solution (*42*) for *ex vivo* glycocalyx permeability assay, 4% PFA for histology and a dry vial frozen at -80°C.

**Mice**

The *Muc17^fl/fl^* mouse strain was engineered by introducing *loxP* sites flanking exons 3 and 5 of the *Muc17* gene (Ensembl gene ID: ENSMUSG00000037390; NCBI gene ID: 666339) (fig. S2A). To generate the *Muc17^∆IEC^* strain; *Muc17^fl/fl^* were crossed with *Vil1-Cre* mice (JAX Strain# 004586, RRID:IMSR_JAX:004586). All mice were on a C57BL/6N background, maintained under standardized SPF conditions of temperature (21–22°C), and illumination (12-hour light/dark cycle) with *ad libitum* access to food and water. Experimental groups consisted of age- and sex-matched 6–8-week-old cohoused littermates, or 5, 8- or 36-40-week-old mice where indicated. Animals were anesthetized with isoflurane followed by cervical dislocation. The Swedish Laboratory Animal Ethical Committee in Gothenburg (ethical permits 2285-19 and 2292-19) approved all experiments.

**Genotyping using PCR**

Ear punch biopsies or 3 mm tissue biopsies from mice were collected and frozen at -20°C. Genomic extraction was performed by addition of 65 µL of alkaline lysis buffer (25 mM NaOH, 0.2 mM EDTA, pH 12) and boiling at 95°C for 1 hour. Samples were neutralized with 65 µL 40 mM Tris-HCl, pH 5. 3 µL of the DNA extraction was used as a template in genotyping PCR reactions using HotStarTaq DNA polymerase (Qiagen Cat# 203205) and 10 µM of each primer. For genotyping of the two *loxP* sites in the *Muc17* gene, forward primer GH979F (5’-CAAACAGTGTCATACCCACTATGG-3’) and reverse primer GH979R (5’-GGCTTTTGATGTTTGATTGTTG-3’), and forward primer GH980F (5’- CCTTAGAGGCATATTGTTCTCAGC-3’) and reverse primer GH980R (5’- TGCCAGCATAAATCAGAGCC-3’) were used (fig. S2A). Thermocycling parameters were denaturation step at 95°C for 5 min., 35 cycles of 95°C for 30 s, 58°C for 50 s, 72°C for 3 min., and final elongation 72°C for 10 min.. For genotyping of the *Vil1-Cre* allele*,* forward primer 1878 (5’- GTGTGGGACAGAGAACAAACC-3’) and reverse primer 1879 (5’- ACATCTTCAGGTTCTGCGGG-3’) were used. Thermocycling parameters were denaturation step 95°C for 5 min., 35 cycles of 95°C for 30 s, 53°C for 50 s, 72°C for 1.5 min., and final elongation 72°C for 10 min..

**Dextran sodium sulfate (DSS)-induced colitis**

Six- to eight-week-old, cohoused female and male *Muc17^fl/fl^* and *Muc17^∆IEC^* littermates we subjected to *ad libitum* administration of 3% DSS (TdB Labs Cat# DB001) in the drinking water. The end point of the experiment was seven days of DSS treatment, loss of 10% of initial weight, or death. Stool consistency, hematochezia (fecal blood), and disease activity scores were calculated for animals that reached day six of treatment. The disease activity index (DAI) was calculated as the sum of the combined scores for stool consistency, fecal blood, and weight loss (*43*). The detection of occult blood was performed using the Hemoccult Guaiac Fecal Occult Blood Test kit (Beckman Coulter. Cat# 61200A) according to the manufacturer’s instructions. The probability of survival was defined by time of death or weight loss >10%. Jejunum and colon were dissected, and the colon length was measured from cecum to anus and normalized against the initial body weight of the respective animal. Segments of the jejunum and distal colon were fixed in Carnoy’s fixative (60% absolute methanol, 30% chloroform, and 10% glacial acetic acid), prepared for sectioning, and stained for hematoxylin/eosin and Alcian Blue-PAS.

**Infection of mice with *Citrobacter rodentium***

Streptomycin-resistant *C. rodentium* strain DBS100 (*44*) and chloramphenicol-resistant DBS100-derived *C. rodentium^GFP+^* (*45*) strains were used in the infection experiments. Infection inocula were cultured overnight in LB broth at 37°C in an orbital shaking incubator. Overnight cultures were concentrated 10-fold by centrifugation at 4000 RCF for 10 min. and resuspension in LB broth. Six- to eight-week-old, cohoused female and male *Muc17^fl/fl^* and *Muc17^∆IEC^* littermates were gavaged with 200 μL of infection inoculum (2 x 10^8^ CFU). The end point of the experiment was as indicated in each experiment, loss of 10% of initial weight, or death. *C. rodentium* load at different tissue sites was determined by sacrificing the mice and collecting samples under aseptic conditions. Approximately 3 cm of jejunum, ileum, and distal colon were dissected and flushed with 4 mL sterile PBS to collect the contents. Contents and flushed tissue were collected separately. *C. rodentium* load was also quantified by sampling the liver, mesenteric lymph nodes, and spleen. Tissue samples were homogenized in sterile PBS using an Ultra-Turrax T10 dispersing instrument (IKA) that was sequentially cleaned in 70% ethanol (×2) and sterile PBS. *C. rodentium* was enumerated from the homogenates by serial dilution on MacConkey agar plates supplemented with 100 µg/mL streptomycin or 30 µg/mL chloramphenicol, followed by overnight incubation at 37°C and quantification of bacterial CFUs. For each sample, a theoretical limit of detection (LOD) was calculated based on the detection of one colony at the lowest plated dilution. An average LOD is shown for each tissue sample.

**Calculation of relative risk**

The relative risk of *C. rodentium* infection based on genotype and tissue was determined by calculating the ratio of the risk of *Muc17^∆IEC^* mice carrying *C. rodentium* counts above LOD to the risk of *Muc17^fl/fl^* mice carrying C. rodentium counts above LOD for each tissue site.

**Tissue histology and immunohistochemistry**

Human ileal biopsies or murine intestinal segments were fixed in Carnoy’s fixative or 4% paraformaldehyde (PFA) solution. Fixed samples were embedded in paraffin, deparaffinized in xylene substitute (2 × 10 min., 60°C), and rehydrated in 100% ethanol (10 min.), 70% (v/v) ethanol (5 min.), 50% (v/v) ethanol (5 min.), and 30% (v/v) ethanol (5 min.). Sections were placed in antigen retrieval buffer (0.01M citric acid, pH 6.0) at 100 degrees for 10 min. and then cooled to RT (2 hours) and transferred to PBS. Tissues were enclosed with a hydrophobic PAP pen, permeabilized with 0.1% Triton X-100 in PBS for 5 min., and then blocked with 5% fetal calf serum (FCS) in PBS for 2 hours at room temperature (RT). Primary (overnight) and secondary antibodies (2 hours) were added to the dilution buffer (5% FCS in PBS). Primary antibodies were rabbit anti-CDHR5 polyclonal antibody (1:250) (Atlas Antibodies Cat# HPA009081, RRID:AB_1079428), rabbit anti-Epcam polyclonal antibody (1:1000) (Abcam Cat# ab71916, RRID: AB_1603782), mouse anti-Ezrin monoclonal antibody (1:500) (Sigma-Aldrich Cat# E8897, RRID: AB_476955), mouse anti-GFP monoclonal antibody (1:500) (Sigma-Aldrich Cat# G6539, RRID: AB_259941), mouse anti-mKi67 monoclonal antibody (1:100) (Thermo Fisher Scientific Cat# 14-5698-82, RRID:AB_10854564), rabbit anti-Muc13 polyclonal antibody against human and mouse MUC13 (*46*) (1:500), rabbit anti-MUC17C1 polyclonal antibody against human MUC17 (*47*) (1:500), and rabbit anti-Muc17S2 polyclonal antibody against mouse Muc17 (*8*) (1:2,000). Secondary antibodies were donkey anti-rabbit IgG Alexa Fluor 488 (1:500) (Thermo Fisher Scientific Cat# A-21206 (also A21206), RRID: AB_2535792), goat anti-mouse IgG1 Alexa Fluor 555 (1:500) (Thermo Fisher Scientific Cat# A-21127, RRID: AB_2535769) and goat anti-rabbit Alexa Fluor 647 (1:500) (Thermo Fisher Scientific Cat# A-21245 (also A21245), RRID: AB_2535813). Brush border glycans were stained with 1 µg/mL biotinylated Aleuria aurantia lectin (AAL) (Vector Laboratories #B-1395-1) or wheat germ agglutinin (WGA) (Vector Laboratories #B-1025-5) overnight and incubated with Streptavidin Alexa Fluor 555 (1:500) (Thermo Fisher Scientific Cat# S21381, RRID:AB_2307336) for 2 hours. DNA was stained with 5 μg/mL Hoechst 34580 (ThermoFisher Scientific Cat# H21486) for 10 min.. Slides were washed three times with PBS after each incubation step. Coverslips were mounted using Prolong Gold antifade (ThermoFisher Scientific Cat# P36980) and polymerized for 2 hours at RT. Slides were imaged using an upright LSM 700 Axio Examiner Z.1 confocal imaging system (Carl Zeiss). AB-PAS stain was performed at pH 2.1-2.5. Assessment of tissue morphology, as measured by crypt number, villus and crypt length, and goblet cell number, in each group was performed in a blinded manner.

**TUNEL assay**

TUNEL assay was performed using In Situ Cell Death Detection Kit, TMR Red (Roche Cat# 12156792910). Rehydrated tissue sections on glass slides were subjected to antigen retrieval and permeabilization as described above. Following the blocking of sections with 3% BSA and 20% bovine serum in PBS for 30 min., slides were rinsed in PBS and sections were incubated with a TUNEL reaction mixture for 1 hour at 37°C in a humidified atmosphere in the dark. Negative control slides were prepared with label solution without terminal transferase. Positive control slides were pretreated with 3U /mL recombinant grade I DNase I in 50 mM Tris-HCl, pH 7.5, 10 mM MgCl_2_, 1 mg/ml BSA for 10 min. at RT before incubation with TUNEL reaction mix. Slides were rinsed three times with PBS and stained with 5 μg/mL Hoechst 34580 for 10 min.. After three washes with PBS, slides were mounted as described above.

**Bioorthogonal labeling of mucin *O*-glycans**

Mice were intraperitoneally injected with 2.6 mg Tetraacetylated N-Azidoacetylgalactosamine (GalNAz) (ThermoFisher Scientific Cat# C33365) dissolved in 25 μL DMSO and diluted in a final volume of 500 µL PBS. Intestinal tissues were collected 8 hours after injection, fixed in 4% PFA, and prepared for paraffin sectioning. Sections were rehydrated as described previously. Click reaction was performed prior to blocking using the Click-IT™ Tetramethylrhodamine (TAMRA) Protein Analysis Detection Kit (ThermoFisher Scientific Cat# C33370) following the manufacturer’s protocol. DNA was stained with 5 μg/mL Hoechst 34580 for 10 min. (ThermoFisher Scientific Cat# H21486).

***Ex vivo* glycocalyx permeability assay in ileal biopsies**

The barrier integrity of the glycocalyx was quantified as previously described (*9*). In detail, biopsies from the ileum of patients were collected in ice-cold oxygenated Krebs transport solution and pinned down with the epithelial side up in silicon-well plates. Mounted samples were stained with 50 μg/mL CellMask Deep Red plasma membrane stain (Thermo Fisher Scientific Cat# C10046) for 15 min. at room temperature (RT), fixed with 4% PFA for 1 hour at RT, and washed three times with PBS. Bacterial penetration into the glycocalyx was quantified using viable *E. coli^GFP+^* (*9*), cultured overnight at 37°C in LB medium supplemented with 100 μg/mL ampicillin and washed in PBS before the assay. Mounted biopsies were incubated with *E. coli^GFP+^* for 20 min. prior to time-lapse confocal imaging (pixel dwell time 2.41 μsec, frame time 20.39 s) for 30 cycles using a Plan-Apochromat × 20/1.0 DIC water-immersion objective (Zeiss), 488/639-nm lasers, on an upright LSM 700 Axio Examiner Z.1 confocal imaging system (Carl Zeiss) with Zen acquisition software (Carl Zeiss). The spatial distance of individual *E. coli^GFP+^* cells relative CellMask was extracted as a function of time using Imaris software (Oxford Instruments). The CellMask-stained brush border was reconstructed using the Isosurface function and shortest distance calculation. Individual *E. coli^GFP+^* cells were reconstructed using the Spot function and shortest distance calculation. The frequency distribution of *E. coli^GFP+^* cells in the distance interval of 0-2 µm from the CellMask-stained brush border was visualized in Prism 10 software (GraphPad). Quantification was performed on an average of 3 regions per villus and three villi per patient biopsy.

***Ex vivo* quantification of mucus barrier properties in murine distal colon**

Quantification of mucus thickness and penetrability in the murine distal colon was performed as previously described (*48*). The distal colon was excised and flushed with ice-cold oxygenated Krebs buffer to remove luminal material, opened along the longitudinal axis, and mounted in a horizontal perfusion chamber (*49*). The tissue was overlaid with Krebs buffer containing a mixture of 5 μg/mL Hoechst 34580 (ThermoFisher Scientific Cat# H21486), 10 µg/mL UEAI Rhodamine (Vector Laboratories Cat# RL-1062-2), and 1 μm crimson carboxylate-modified FluoSpheres microbeads (1:20 dilution) (Thermo Fisher Scientific Cat# F8816) and incubated for 15 min.. The tissue was then washed with 0.5 mL Krebs buffer and submerged in 2 mL fresh Krebs buffer. Tissue incubated with microbeads was imaged using the 488/546/639-nm lasers on an LSM 700 Axio Examiner Z.1 confocal imaging system (Carl Zeiss) as described above. The colonic epithelium (Hoechst), mucus (UEAI), and microbead fluorescent signals were mapped using Imaris software (Oxford Instruments) and data describing the z-axis position of epithelium and microbeads was extracted. The thickness of the mucus layer was quantified by calculating the average epithelium-microbead z-axis distance. Normalized mucus penetrability was quantified by quantifying the distribution of microbeads within the mucus layer stained with UEAI. A frequency distribution curve of the axis distance of microbeads from the epithelium was generated for each z stack using Prism 10 software. Curves were normalized to maximum frequency values and then normalized to the position of the mucus surface and cropped to exclude data from microbeads above the mucus surface. Lastly, normalized penetrability was expressed as the area under the curve to allow for a quantitative comparison of microbead penetration between samples.

**Isolation of luminal vesicles from intestinal segments**

Isolation of luminal vesicles was adapted from (*50*). Briefly, mice were sacrificed as described above and the intestines were transferred to ice-cold saline (150 mM NaCl, 2 mM imidazole-Cl) containing 0.02% NaN_3_. The intestinal content was collected in a glass beaker by flushing the intestine with 30-60 mL of ice-cold saline. The content was centrifuged at 500 x g for 20 min., 20,000 x g for 30 min., and finally 100,000 x g for 2 hours. The supernatant from the proceeding centrifugation was used in the subsequent step. All centrifugation steps were conducted at 4°C. The final pellet, composed of luminal vesicles, was resuspended in PBS.

**Electrophoresis and western blot**

A suspension of luminal vesicles was reduced in 4X reducing sample buffer (8% SDS, 400 mM Dithiothreitol) and separated on precast 4%–12% SDS-polyacrylamide gel (ThermoFisher Scientific Cat# XP04125BOX). Proteins were transferred to a PVDF-FL membrane (Millipore Cat# IPFL00010) with a current of 2.5 mA/cm2 for 1 hour. The membrane was blocked in 5% non-fat milk in PBS for 30 min. and incubated with rabbit anti-Muc17C1 polyclonal antibody against mouse Muc17 (*51*) (1:250) in 5% non-fat milk in PBS + 0.1% Tween-20 (PBS-T) overnight at 4°C. The membrane was washed three times in PBS-T and incubated with Goat anti-rabbit Alexa Fluor 680 (1:20000) (Thermo Fisher Scientific Cat# A-21109, RRID:AB_2535758) secondary antibody for 1 hour at RT in the dark. The membrane was washed three times in PBS-T and visualized on an Odyssey CLx near-infrared fluorescence imaging system (LI-COR Biosciences).

**16S rRNA gene sequencing for profiling of intestinal microbiota**

DNA from luminal compartments was extracted by mechanical lysis using a Fast-Prep System with Lysing Matrix E tubes (MPBio) as previously described (*52*). Sample concentrations were measured using a Qubit instrument (Thermo Fisher Scientific Cat# Q33238) with the Qubit™ 1X dsDNA High Sensitivity (HS) (Thermo Fisher Scientific Cat# Q33230). Luminal microbiota composition was profiled by sequencing the V3/V4 region of the 16S rRNA gene using a 2-step PCR approach (*53*). The forward and reverse primer mixes were prepared by combining eight phased primers each, at equimolar levels. For primer sequences see table S3. The PCR1 reaction was performed with 1 ng of input DNA, 10 µl KAPA HiFi Hotstart 2x Master Mix (Roche Cat# KK2602), 0.5 µl BSA (20 mg/ml), 1 µl of 7.5 µM forward primer mix and 1 µl of 7.5 µM reverse primer mix. Nuclease-free water was added to a reaction size of 21 µl. The PCR conditions for PCR1 were 98°C for 2 min., 20 cycles of 98°C for 20 s, 54°C for 20 s, 72°C for 15 s, and final elongation at 72°C for 2 min.. Samples were purified using 21 µl of MagSI NGSPrep plus purification beads (Magtivio Cat# MDKT0001) following the supplier’s instructions and samples were eluted in 12 µl of Elution Buffer. 6 µl of the PCR1 eluate was subjected to amplification (PCR2 reaction) with the following reaction setup: 10 µl KAPA HiFi Hotstart 2x Master Mix, 1 µl of 5 µM Adapterama i7 index primer and 1 µl of 5 µM Adapterama i5 index primer. For primer sequences see table S3. Nuclease-free water was added to a reaction size of 20 µl. The PCR conditions for PCR2 were 98°C for 2 min., 8 cycles of 98°C for 20 s, 55°C for 30 s, 72°C for 30 s, and final elongation at 72°C for 2 min.. Samples were purified using 20 µl of MagSI NGSPrep plus purification beads. Libraries were pooled at equimolar levels and the molarity was determined by qPCR using the Illumina library Quantification kit from (Roche Cat# KK4824). Pools were run on a MiSeq V3 2×300 with a loading concentration of 11 pM. PhiX spike-in was 10%. The same library pool of phased libraries was used for all sequencing runs.

The 16S rRNA analysis was performed using the nf-core/ampliseq analysis pipeline (*54*). Bioinformatic analysis was performed with QIIME 2 2020.11 (*55*). Raw sequence data were demultiplexed and quality filtered followed by denoising with DADA2 (*56*). All amplicon sequence variants (ASVs) were aligned with mafft v.7.407 (*57*) and used to construct a phylogeny with fastTree v.2.1.10 (*58*). Alpha-diversity metrics (Shannon diversity index), beta diversity metrics (Bray-Curtis dissimilarity), and Principal Coordinate Analysis (PCoA) were estimated using the diversity core-metrics-phylogenetic command. Taxonomy was assigned to ASVs using the q2-feature-classifier (*59*) classify-Sklearn naïve Bayes taxonomy classifier against the Silva v.138 reference sequence database (*60*). LEfSe algorithm was used to correlate genus-level relative abundance data to different experimental groups (*61*). Microbial features in the luminal compartment of each genotype were correlated to the calculated z-scores representing the relative abundance of each taxon along the gastrointestinal tracts reported by Lkhagva et al. (*27*).

**Quantification of bacteria by 16S rRNA gene qPCR**

For assessment of bacterial density in peripheral tissues, tissues were dissected and flushed with 5 mL 0.22 μm filter-sterilized PBS. Tissues were lysed by brief homogenization using an Ultra-Turrax T10 dispersing instrument (IKA) that was cleaned in RBS detergent (Merck), 70% ethanol, and filter-sterilized ddH2O between each sample. Tissue lysates were centrifuged at 10,000 RCF for 10 min. to pellet bacterial cells and tissue debris, and DNA was extracted using a QIAmp PowerFecal Pro kit (Qiagen Cat# 51804) with 4x rounds of 4.5 m/s for 40 s bead-beating using a Fast-Prep System (MPBio). Each sample was assessed in triplicates by analyzing 45 ng of isolated DNA with qPCR using SsoFast EvaGreen Supermix (Bio-Rad Cat# 1725204) with 0.3 μM universal primers 926F (5’-AAACTCAAAKGAATTGACGG-3’) and 1062R (5’-CTCACRRCACGAGCTGAC-3’) on a CFX96 platform (Bio-Rad). The absolute bacterial 16S copy number was quantified using standard curves generated from qPCR of whole 16S gene amplicons purified from *E. coli*. Data was normalized to the weight of the initial sample.

**Fluorescence *in situ* hybridization**

FISH staining for the bacterial 16S rRNA was performed on tissue sections from Si5 and DC. Sections were deparaffinized, air dried, and incubated with hybridization buffer (40% vol/vol formamide, 0.1% wt/vol SDS, 0.9 M NaCl, and 20 mM Tris, pH 7.4) supplemented with 1 mM Alexa 555-labeled universal bacterial FISH probe EUB338 (*62*). Slides were incubated at 37°C overnight in a RapidFISH Slide Hybridization Oven (Boekel Scientific), rinsed in wash buffer (0.9 M NaCl and 25 mM Tris, pH 7.4), and incubated for 20 min. at 50°C. Finally, slides were rinsed in double-distilled water and counterstained with 5 μg/mL Hoechst DNA stain and Wheat Germ Agglutinin Fluorescein (WGA) (10 µg/mL) (Vector Laboratories Cat# FL-1021) for 15 min. before the mounting of coverslips. Slides were imaged with an LSM700 confocal microscope (Zeiss).

**Plate culture and identification of viable bacteria in extraintestinal tissues**

Liver, mesenteric lymph nodes, and spleen were harvested into 1 mL sterile PBS and homogenized by a sterilized 5 mm stainless steel bead for 40 sec using a Fast-Prep 24 instrument (MP Biomedicals). 100 μL homogenate of each tissue was spread on brain heart infusion supplemented (BHIS) agar plates and incubated at 37˚C under anaerobic conditions for 5 days. Colonies were transferred to 100 µL ddH2O and the bacterial 16S rRNA gene was amplified with the primer 27F (5’-AGAGTTTGATYMTGGCTCAG-3’) and 1492R (5’-GGTTACCTTGTTACGACTT-3’) using *PfuUltra* High-Fidelity DNA Polymerase (Agilent Cat# #600380). Thermocycling conditions were initial denaturation step of 95°C for 1 min., 35 cycles of 95°C for 30 s, 48°C for 30 s and 72°C for 2 min., and an extension step 72°C for 10 min.. Amplicons were separated on 1.5% agarose gel, excised and purified using a Nucleospin PCR Clean Up kit (Macherey Nagel Cat# 740609.50) and subjected to DNA sequencing (Eurofins Genomics) using sequencing primers targeting the hypervariable region V3-V4 of the 16S rRNA gene, S-D-Bact-0341-b-S-17(5’-CCTACGGGNGGCWGCAG-3’) and the S-D-Bact-0785-a-A-21 (5’-GACTACHVGGGTATCTAATCC-3’) (*63*). 16S gene sequences were analyzed by the BLASTn tool (NCBI) to identify bacterial isolates.

**Single-cell analysis of published data sets**

Single-cell sequencing data of the human small intestine and colon were extracted from GSE185224 (*64*) using the Gene Expression tool to query CZ CELLxGENE Discover (Chan Zuckerberg Initiative). Single-cell sequencing data of the mouse small intestine was obtained from GSE92332 (*65*) and extracted from the Single Cell Portal (Broad Institute).

**Proteomic profiling of intestinal epithelial cells**

Jejunal segments were opened longitudinally, washed in PBS for 5 min., and then incubated in PBS containing 3 mm EDTA and 1 mm DTT at 4 °C for 1 hour while gently shaken. The solution was replaced with fresh PBS, and epithelial cells were dissociated from the tissue by vigorous shaking for 30 s. The remaining tissue was removed from the solution using forceps, and cells were pelleted via centrifugation at 500 rpm. Isolated epithelial cells were lysed in 400 μL lysis buffer (100 mM DTT, 4% SDS, 100 mM Tris pH 7.5) and heated at 95°C for 5 min.. Lysates were sonicated for 10 s, centrifuged at 16,000 x g at RT and the supernatant was added onto 10 kDa cutoff filters (PALL Cat# OD010C33). Proteins were digested using filter-aided sample preparation (*66*) with trypsin at 37°C overnight. Peptide concentration after elution was measured at 280 nm using NanoDrop (Thermo Fisher Scientific) and peptides were cleaned with StageTip C18 columns (*67*) before mass spectrometry (MS).

Nano LC-MS/MS was performed on a Q-Exactive HF mass-spectrometer (Thermo Fischer Scientific), connected with an EASY-nLC 1000 system (Thermo Fischer Scientific) through a nanoelectrospray ion source. Peptides were loaded on a reverse-phase column (150 mm x 0.075 mm inner diameter, New Objective, New Objective, Woburn, MA), packed in-house with Reprosil-Pur C18-AQ 3 mm particles (Dr. Maisch, Ammerbuch, Germany). Peptides were separated with a 230-minute gradient: from 3% to 25% B in 175 min., 25% to 45% B in 30 min., 45% to 100% B in 5 min., followed 20 min. wash with 100% of B (A: 0.1% formic acid, B: 0.1% formic acid/80% acetonitrile) using a flow rate of 250 nl/min.. Q-Exactive HF was operated at 250°C capillary temperature and 2.0 kV spray voltage. Full mass spectra were acquired in the Orbitrap mass analyzer over a mass range from m/z 350 to 1600 with a resolution of 60 000 (m/z 200) after the accumulation of ions to a 3e6 target value based on predictive AGC from the previous full scan. The twelve most intense peaks with a charge state ≥ 2 were fragmented in the HCD collision cell with a normalized collision energy of 27%. The tandem mass spectrum was acquired in the Orbitrap mass analyzer with the resolution of 15,000 after accumulation of ions to a 1e5 target value. Dynamic exclusion was set to 30 s. The maximum allowed ion accumulation times were 20 ms for full MS scans and 50 ms for tandem mass spectrum.

MS raw files were analyzed with MaxQuant software version 1.5.7.4 (*68*). Peak lists were identified by searching against the mouse UniProt protein database (downloaded 2022.11.11), supplemented with an in-house database containing all the mouse mucin sequences (http://www.medkem.gu.se/mucinbiology/databases/). Searches were performed using trypsin as an enzyme, maximum 2 missed cleavages, and a precursor tolerance of 20 ppm in the first search used for recalibration, followed by 7 ppm for the main search and 0.5 Da for fragment ions. Carbamidomethylation of cysteine was set as a fixed modification, and methionine oxidation and protein N-terminal acetylation were set as variable modifications. The required false discovery rate (FDR) was set to 1% for peptide and protein levels, and the minimum required peptide length was set to seven amino acids. Label-free quantification (LFQ) was based on two peptides. Raw mass spectrometry data were analyzed with DEP package (*69*) for R. Briefly, intensities were background corrected and normalized by variance stabilizing transformation and filtered based on valid values criteria (a minimum of 3 valid values in at least one of the groups). Missing values were imputed with a width of 0.3 of the Gaussian distribution relative to the standard deviation of measured values and a downshift of 1.8 units of the standard deviation of the valid data. Volcano plots were generated to visualize differentially expressed proteins between two conditions.

**Statistical Analysis**

Statistical analysis and graphical illustrations were performed using Prism 10 software (GraphPad) and R open-source software. Biological replicates for each group and experiment are stated in the figure legends. All data are presented as means ± SD except the quantification CFU of *C. rodentium* (Fig. 4D-F, fig. S5) as medians with interquartile range, quantification of total weight as median with minimum to maximum range of data points (Fig. 8A), and the quantification of the 16S rRNA gene in peripheral tissues (Fig. 8C), which is represented as mean ± SEM. Statistical tests were applied as indicated in each figure legend and the p-values for each statistical test are indicated in each. Principal components analysis (PCA) of genotype- and age-dependent epithelial and mucus parameters was performed using the ‘princomp’ function in R.

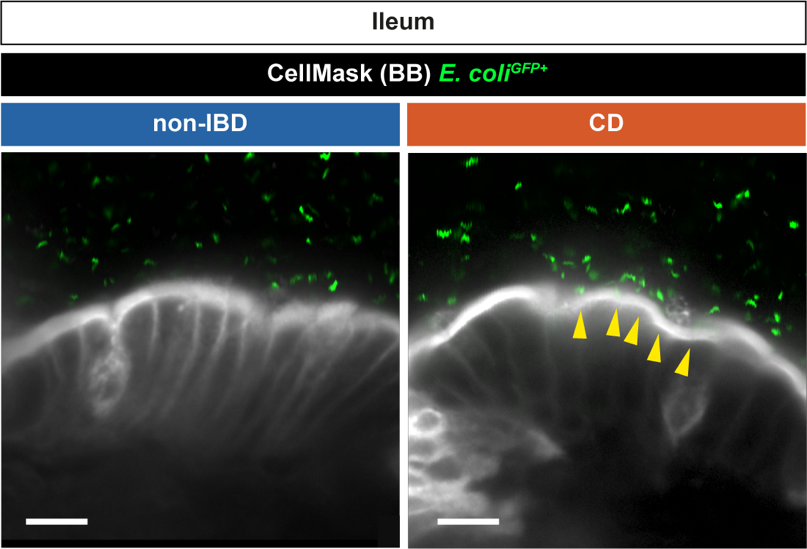

**Fig. S1. Confocal micrographs of biopsies from non-IBD and CD patients analyzed using *ex vivo* glycocalyx permeability assay.**

Representative confocal micrographs of biopsy explants, stained for the brush border (CellMask, white) and incubated with *E. coli^GFP+^* (green). Yellow arrows indicate points of contact between *E. coli^GFP+^* and the brush border. Scale bar 10 µm.

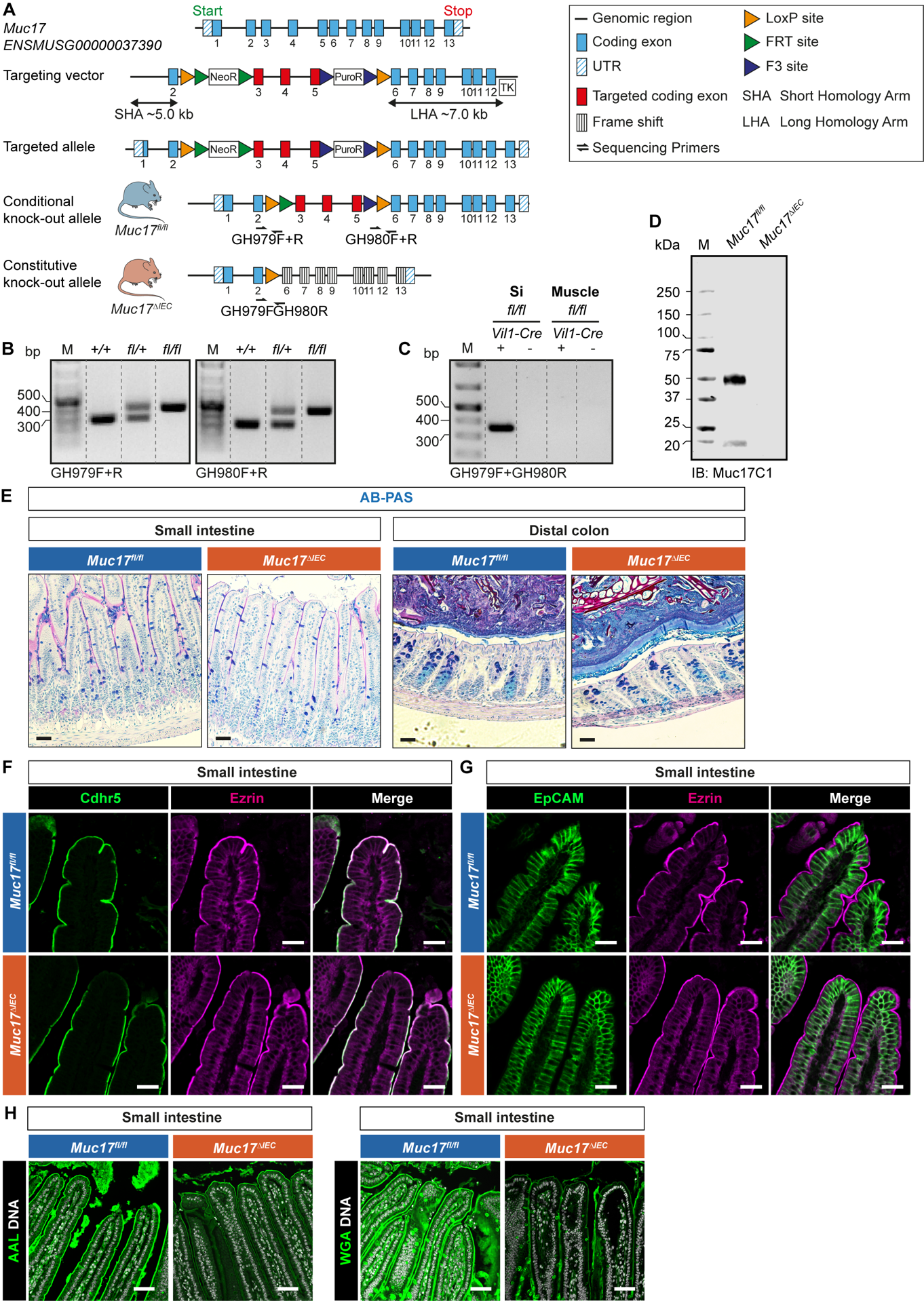

**Fig. S2. Characterization of the *Muc17^∆IEC^* mouse strain.**

**(A)** Schematic illustration of the murine *Muc17* gene and the strategy for introducing *loxP* sites after the exons 2 and 5. The binding sites for primers GH979F+R and GH980F+R are indicated with black arrows. **(B)** Representative DNA agarose gels showing genotyping of the floxed *Muc17* allele. M, molecular weight. **(C)** Representative DNA agarose gels showing genotyping of the floxed *Muc17* alleles in the presence and absence of the *Vil1-Cre* allele in the small intestine (Si) and muscle tissues. M, molecular weight. **(D)** Immunoblot (IB) of luminal vesicles isolated from the small intestine of *Muc17^fl/fl^* and *Muc17^∆IEC^* mice. **(E)** AB-PAS staining of the jejunum (Si5) and distal colon (DC) of *Muc17^fl/fl^* and *Muc17^∆IEC^* mice. Scale bar 50 µm. **(F)** Immunohistochemistry of Cdhr5 (green) and Ezrin (magenta) in histological sections of the small intestine in *Muc17^fl/fl^* and *Muc17^∆IEC^* mice. Scale bar 20 µm. (**G**) Immunohistochemistry of Epcam (green) and Ezrin (magenta) in histological sections of the small intestine in *Muc17^fl/fl^* and *Muc17^∆IEC^* mice. Scale bar 20 µm. (**H**) Immunohistochemistry using the lectins AAL and WGA in the small intestine of *Muc17^fl/fl^* and *Muc17^∆IEC^* mice. Scale bar 50 µm.

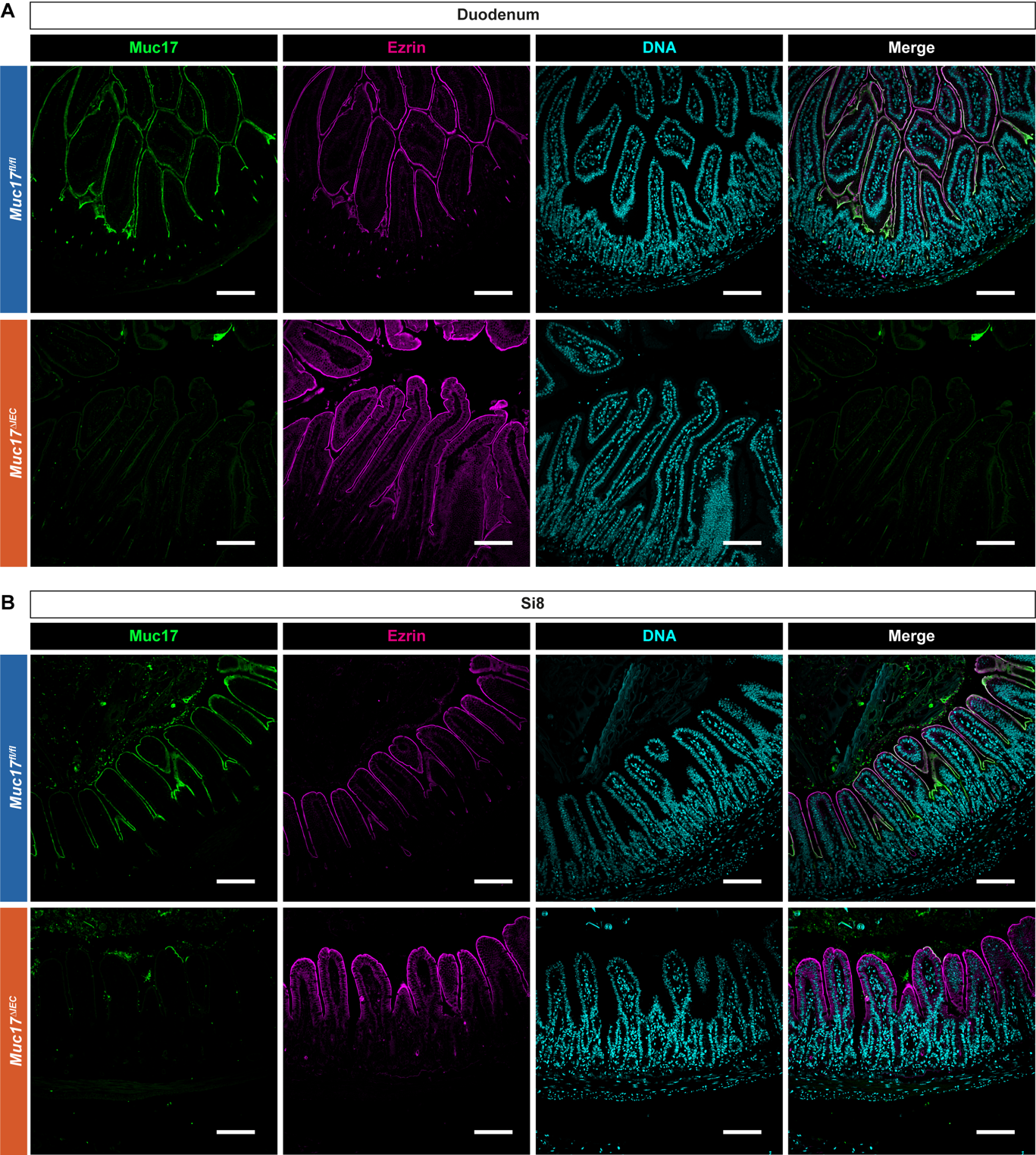

**Fig. S3. Confocal micrographs of the small intestine of *Muc17^fl/fl^* and *Muc17^∆IEC^* mice.**

**(A)** Immunohistochemistry of Muc17 (green), Ezrin (magenta) and DNA (cyan) in histological sections from the duodenum of *Muc17^fl/fl^* and *Muc17^∆IEC^* mice. Scale bar 100 µm. **(B)** Immunohistochemistry of Muc17 (green), Ezrin (magenta) and DNA (cyan) in histological sections from the ileum (Si8) of *Muc17^fl/fl^* and *Muc17^∆IEC^* mice. Scale bar 100 µm.

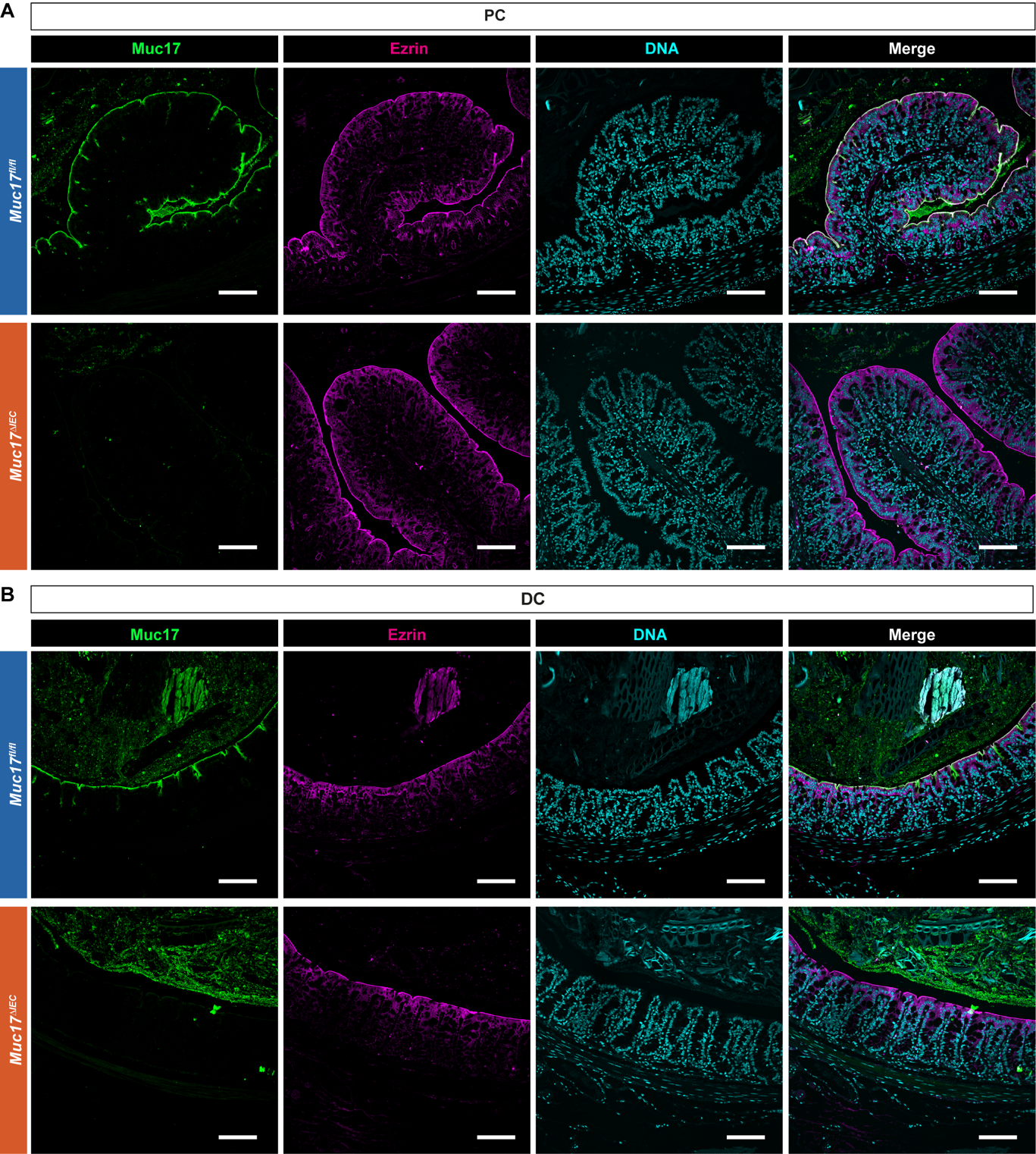

**Fig. S4. Confocal micrograph of the colon of *Muc17^fl/fl^* and *Muc17^∆IEC^* mice.**

**(A)** Immunohistochemistry of Muc17 (green), Ezrin (magenta) and DNA (cyan) in histological sections from the proximal colon (PC) of *Muc17^fl/fl^* and *Muc17^∆IEC^* mice. Scale bar 100 µm. **(B)** Immunohistochemistry of Muc17 (green), Ezrin (magenta) and DNA (cyan) in histological sections from the distal colon (DC) of *Muc17^fl/fl^* and *Muc17^∆IEC^* mice. Scale bar 100 µm.

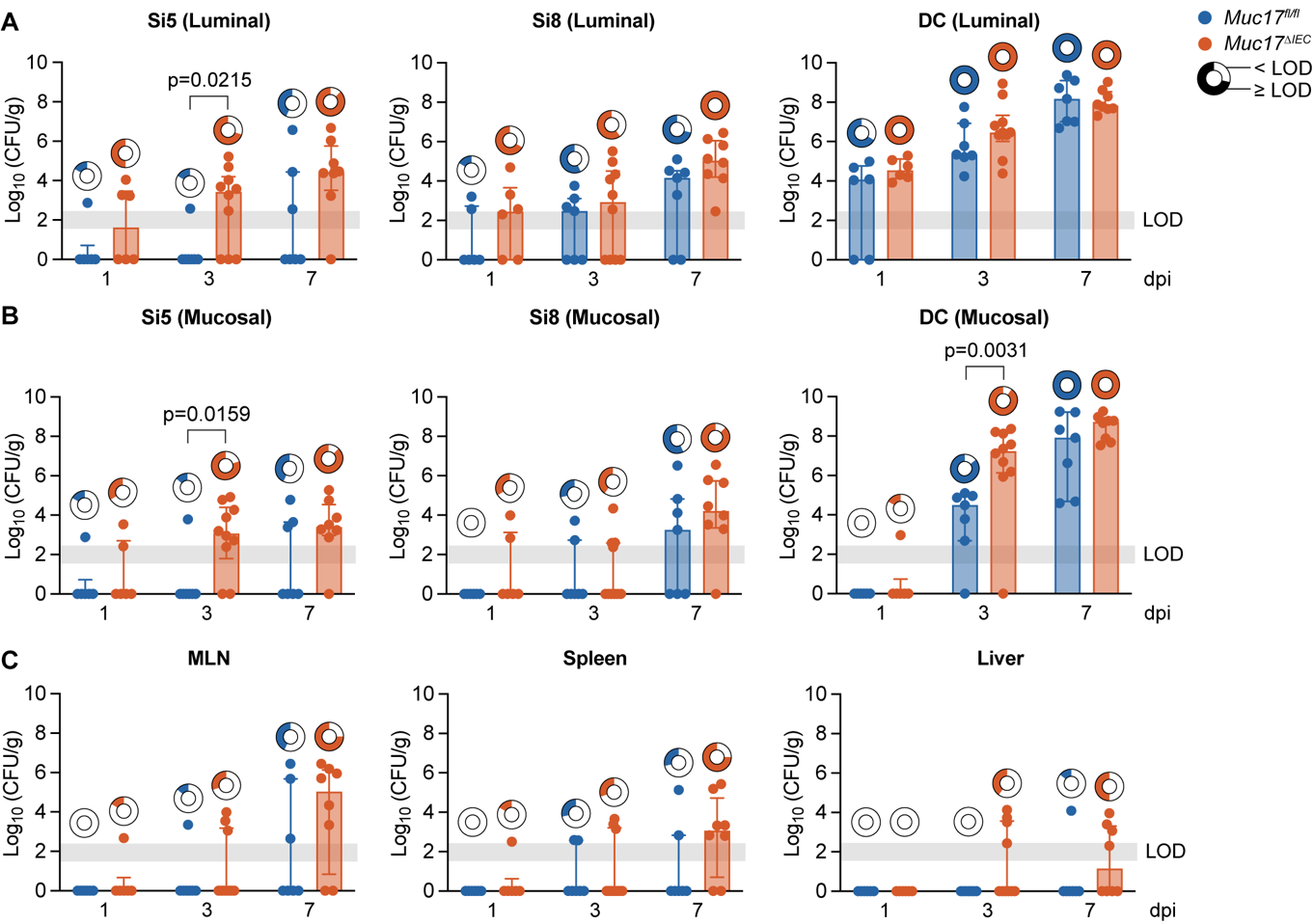

**Fig. S5. Longitudinal analysis of *C. rodentium* infection in *Muc17^fl/fl^* and *Muc17^∆IEC^* mice.**

**(A)** Quantification of *C. rodentium* colony forming unites (CFU) in the luminal compartment of the jejunum (Si5), ileum (Si8), and distal colon (DC) of *Muc17^fl/fl^* and *Muc17^∆IEC^* mice at 1, 3, and 7 days post-infection (dpi). **(B)** Quantification of *C. rodentium* CFU in the mucosal compartment of Si5, Si8, and DC of *Muc17^fl/fl^* and *Muc17^∆IEC^* mice at 1, 3, and 7 dpi. (**C**) Quantification of *C. rodentium* CFU in the mesenteric lymph nodes (MLN), spleen, and liver of *Muc17^fl/fl^* and *Muc17^∆IEC^* mice at 1, 3, and 7 dpi. The circle charts represent the proportion of mice carrying *C. rodentium* CFU above the limit of detection (LOD) in each segment of each genotype. Significance was determined by Mann-Whitney test (A-B).

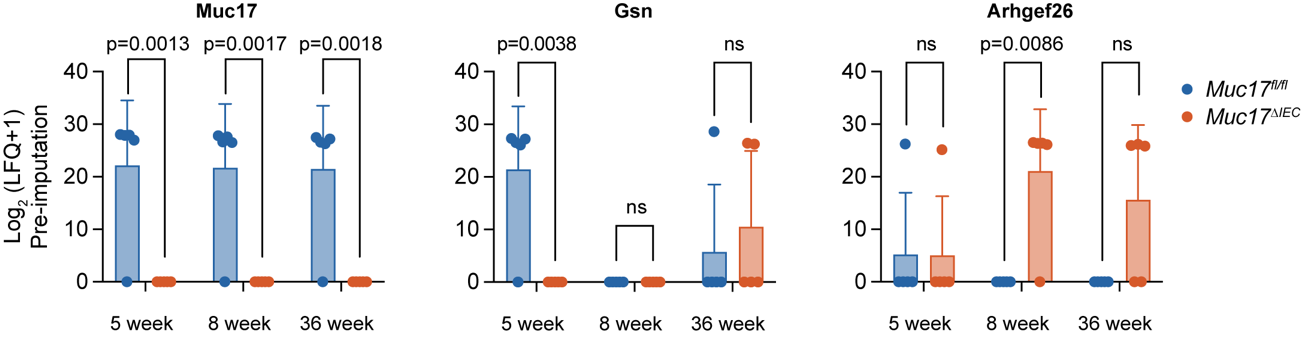

**Fig. S6. Longitudinal proteomic analysis of isolated jejunal epithelial cells from *Muc17^fl/fl^* and *Muc17^∆IEC^* mice.**

Abundance of selected proteins in 5-, 8-, and 36-week-old *Muc17^fl/fl^* and *Muc17^∆IEC^* mice. Significance was determined by two-way ANOVA followed by Šidák correction.

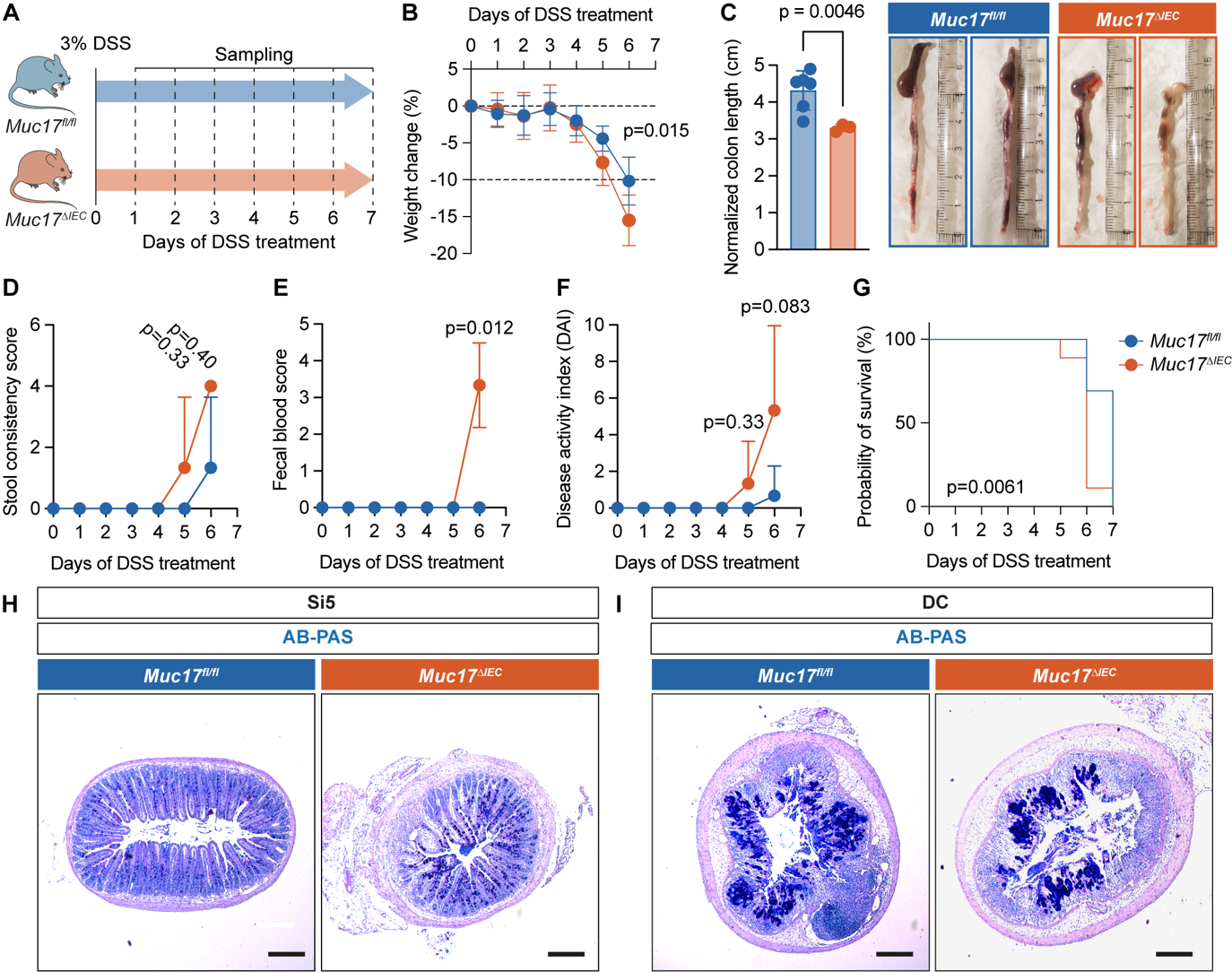
**Fig. S7. Muc17 does not protect against experimental colitis.**

(**A**) Schematic illustration of the DSS challenge protocol and sampling time points. (**B**) Weight loss of DSS-treated *Muc17^fl/fl^* and *Muc17^∆IEC^* mice, depicted as percent of the initial weight at the start of the challenge (time point 0 days). n=13 for *Muc17^fl/fl^*, n=9 for *Muc17^∆IEC^*. (**C**) Weight-normalized colon length. n=6 for *Muc17^fl/fl^*, n=3 for *Muc17^∆IEC^*. Representative images of colonic segments from *Muc17^fl/fl^* and *Muc17^∆IEC^* mice exposed to the DSS challenge. (**D**) Stool consistency score of *Muc17^fl/fl^* and *Muc17^∆IEC^* mice after the DSS challenge. n=6 for *Muc17^fl/fl^*, n=3 for *Muc17^∆IEC^*. (**E**) Fecal blood score of *Muc17^fl/fl^* and *Muc17^∆IEC^* mice after DSS challenge. n=6 for *Muc17^fl/fl^*, n=3 for *Muc17^∆IEC^*. (**F**) Disease activity index of *Muc17^fl/fl^* and *Muc17^∆IEC^* mice after DSS challenge. n=6 for *Muc17^fl/fl^*, n=3 for *Muc17^∆IEC^*. (**G**) Probability of survival of *Muc17^fl/fl^* and *Muc17^∆IEC^* mice exposed to DSS challenge. n=13 for *Muc17^fl/fl^*, n=9 for *Muc17^∆IEC^*. (**H**) Histological sections of *Muc17^fl/fl^* and *Muc17^∆IEC^* jejunum (Si5) after DSS treatment stained with AP-PAS. Scale bar 100 µm. (**I**) Histological sections of *Muc17^fl/fl^* and *Muc17^∆IEC^* distal colon (DC) after DSS treatment stained with AP-PAS. Scale bar 100 µm. Significance was determined by two-way ANOVA followed by Šidák correction (B), Mann-Whitney test (C-F), and Log-rank (Mantel-Cox) test (G).

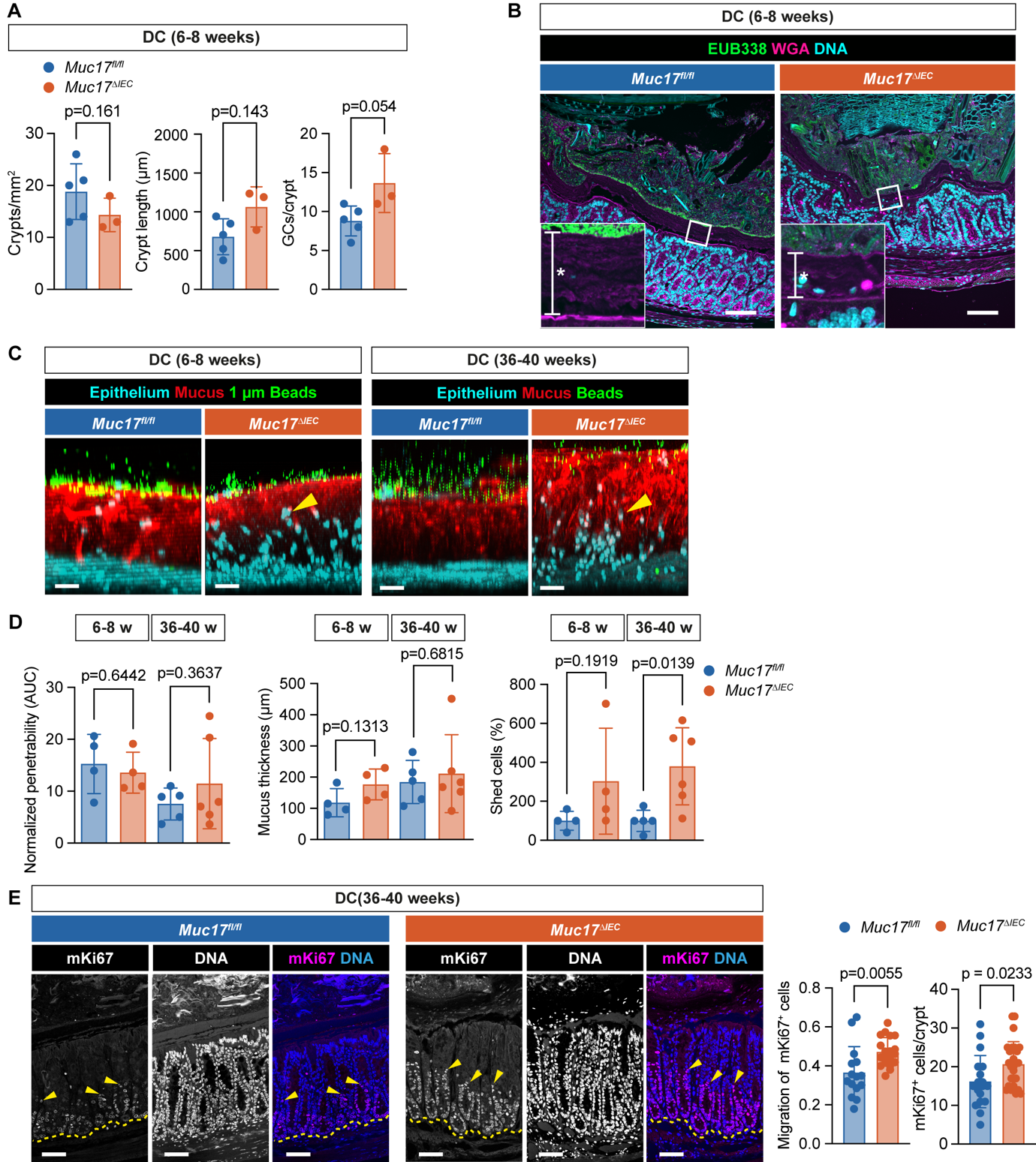
**Fig. S8. Age-dependent mucus barrier defects in the distal colon of *Muc17^∆IEC^* mice housed under baseline SFP conditions.**

(**A**) Quantification of the number of crypts per mm^2^ of histological section, crypt length, and goblet cells (GCs) per crypt in the distal colon (DC) of *Muc17^fl/fl^* and *Muc17^∆IEC^* mice. n=6 for *Muc17^fl/fl^*, n=3 for *Muc17^∆IEC^*. (**B**) Confocal micrographs of *Muc17^fl/fl^* and *Muc17^∆IEC^* DC stained for bacteria (EUB338, green), epithelium and mucus (WGA, magenta), and DNA (cyan). Asterisks mark the inner mucus layer (IML). Scale bar 50 µm. (**C**) Representative images of *ex vivo* mucus penetrability assay in 6–8- and 36–40-week-old *Muc17^fl/fl^* and *Muc17^∆IEC^* DC. DNA (epithelium, cyan), mucus (red), and 1 µm FluoSpheres microbeads (green). Scale bar 30 µm. (**D**) Quantification of mucus height, mucus penetrability, and shed epithelial cells in 6–8- and 36–40-week-old *Muc17^fl/fl^* and *Muc17^∆IEC^* DC. n=4-6 per group. (**E**) Immunohistochemistry of mKi67 (magenta) and DNA (blue) in 36–40-week-old *Muc17^fl/fl^* and *Muc17^∆IEC^* DC. Each channel is also shown in grayscale. Yellow arrows point to mKi67^+^ cells with maximum migration along the crypt axis. The dashed yellow line depicts the crypt bottom. Scale bar 50 µm. Quantification of the migration of mKi67^+^ cells along the crypts and absolute numbers of mKi67^+^ cells in 36–40-week-old *Muc17^fl/fl^* and *Muc17^∆IEC^* DC. n=5-12 crypts per mouse, 3 mice per group. Significance was determined by Mann-Whitney test (A) and unpaired t-test (D-E).

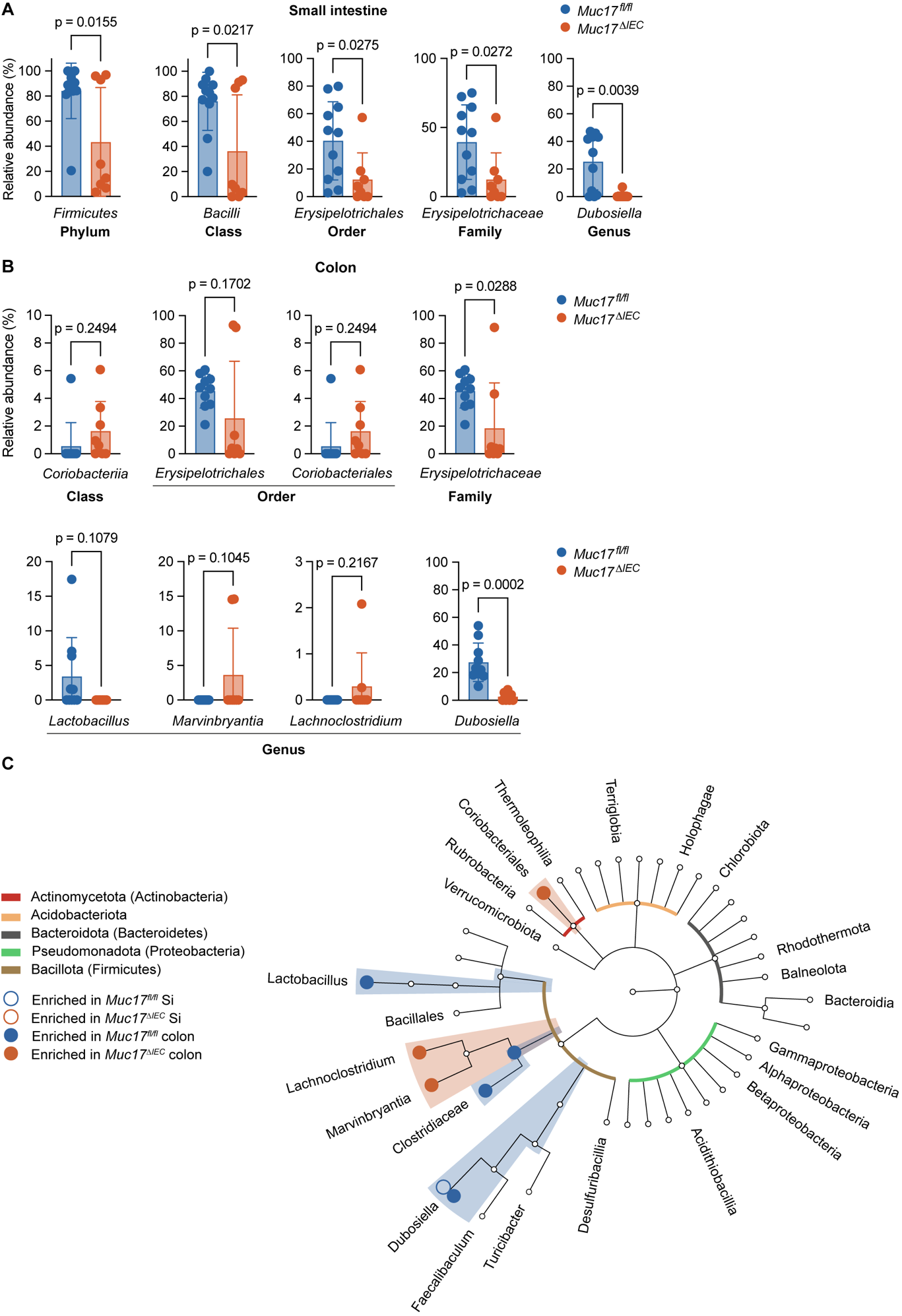
**Fig. S9. Relative abundances of bacterial taxa enriched in *Muc17^fl/fl^* and *Muc17^∆IEC^* mice. (A)** Relative abundance of enriched taxa in the small intestine of *Muc17^fl/fl^* and *Muc17^∆IEC^* mice. **(B)** Relative abundance of enriched taxa in the colon of *Muc17^fl/fl^* and *Muc17^∆IEC^* mice. Significance was determined using unpaired t-test. **(C)** Phylogenetic tree displaying bacterial taxa that are enriched in the small intestine (Si) and colon of *Muc17^fl/fl^* and *Muc17^∆IEC^* mice.

**Table S1. Clinical metadata and sample information. (separate file)**

**Table S2. Longitudinal proteomic profiling of Si5 IECs in *Muc17^fl/fl^* and *Muc17^∆IEC^* mice. (separate file)**

**Table S3. Primer sequences for 16S V3/V4 amplicon library preparation for luminal microbiota profiling and Index sequences for Adapterama scheme. (separate file)**

**Table S4. Primary data for all the described experiments. (separate file)**

**Movie S1. Time-lapse *ex vivo* glycocalyx permeability assay in non-inflamed ileal biopsies from non-IBD and CD patients.**

Representative movies of biopsy explants, stained for the brush border (CellMask, white) and incubated with *E. coli^GFP+^* (green). Yellow arrows indicate points of contact between *E. coli^GFP+^* and the brush border. Scale bar 10 µm.
